# A comparison of humans and baboons suggests germline mutation rates do not track cell divisions

**DOI:** 10.1101/844910

**Authors:** Felix L. Wu, Alva Strand, Carole Ober, Jeffrey D. Wall, Priya Moorjani, Molly Przeworski

**Author notes:** co-supervised this work (Priya Moorjani, Molly Przeworski).

## Abstract

In humans, most germline mutations are inherited from the father. This observation is widely interpreted as resulting from the replication errors that accrue during spermatogenesis. If so, the male bias in mutation should be substantially lower in a closely related species with similar rates of spermatogonial stem cell divisions but a shorter mean age of reproduction. To test this hypothesis, we resequenced two 3–4 generation nuclear families (totaling 29 individuals) of olive baboons (*Papio anubis*), who reproduce at ~10 years of age on average. We inferred sex-specific mutation rates by analyzing the data in parallel with three three-generation human pedigrees (26 individuals). The mutation rate per generation in baboons is 0.55×10^−8^ per base pair, approximately half that of humans. Strikingly, however, the degree of male mutation bias is approximately 3:1, similar to that of humans; in fact, a similar male bias is seen across mammals that reproduce months, years or decades after birth. These results echo findings in humans that the male bias is stable with parental ages and cast further doubt on the assumption that germline mutations track cell divisions. Our mutation rate estimates for baboons raise a further puzzle in suggesting a divergence time between apes and Old World Monkeys of 67 My, too old to be consistent with the fossil record; reconciling them now requires not only a slowdown of the mutation rate per generation in humans but also in baboons.

## Introduction

Germline mutations are the source of heritable differences among individuals. They arise from accidental changes to the genome that occur in the development of a future parent, in the cell lineage from zygote to gamete (egg or sperm), either as errors in replication or due to damage that is improperly corrected by the next round of DNA replication. The number of de novo mutations (DNMs) that arise in an offspring is thus an aggregate over the outcomes of mutational processes that play out from zygote to primordial germ cell and across the cell states of paternal and maternal gametogenesis.

Interestingly, the rate at which DNMs are introduced each generation varies substantially among species: as an illustration, across vertebrates surveyed to date, per base pair (bp) per generation mutation rates span an order of magnitude [summarized in (1), Table 1]. Analyses of DNM and polymorphism data further indicate that in addition to the total rate, the proportion of each mutation type (the “mutation spectrum”) also varies among mammalian species (2–4) and even across human populations (5–8).

**Table 1:**

| Species | Common name | Mean paternal age at conception | Sex-averaged mutation rate per generation, as reported ($\times 10^{-8}$) | Point estimate of $\alpha$, ratio of paternal to maternal DNMs | Number of trios | References |
| --- | --- | --- | --- | --- | --- | --- |
| *Papio anubis* | Baboon | 10.27 | 0.55 | 3.23 | 12 | this study |
| *Homo sapiens* | Human | 34.29 | 1.23 | 3.87 | 10 | this study |
| *Homo sapiens* (Jónsson et al.) | Human | 31.63 | 1.29<sup>A</sup> | 4.05 | 225 | (16) |
| *Pan troglodytes* (Besenbacher et al.) | Chimpanzee | 19.27 | 1.26<sup>B</sup> | 4.37 | 7 | (23,48) |
| *Pan troglodytes* (Tatsumoto et al.) | Chimpanzee | 24.00 | 1.48 | 3.08 | 1 | (47) |
| *Gorilla gorilla* | Gorilla | 13.50 | 1.13<sup>B</sup> | 2.00 | 2 | (23) |
| *Pongo abelii* | Orangutan | 31.00 | 1.66 | 4.13 | 1 | (23) |
| *Macaca mulatta* | Macaque | 7.50 | 0.37 | 1.94 | 14 | (24) |
| *Aotus nancymae* | Owl Monkey | 5.55 | 0.81 | 2.09 | 14 | (22) |
| *Mus musculus* | Mouse | 0.44 | 0.39 | 2.76 | 40 | (4) |
| *Bos taurus* | Cattle | 5.00<sup>C</sup> | 1.2 | 2.53 | 13 | (2,49,50) |

504 <sup>A</sup>Includes single base pair indels505 <sup>B</sup>Mean of reported rates per individual506 <sup>C</sup>Typical age at reproduction drawn from literature. All other ages are as reported in the relevant study.

The composite nature of the per generation mutation rate allows for many possible explanations for these observations. At the cellular level, the damage rates to which species are exposed may differ, and repair and replication efficiencies may change over time. The overall cellular composition of the germline may also evolve—for example, the number of replications per generation. Shifts in any of these components can alter the overall mutation rate. These changes could occur by chance, by genetic drift, or due to natural selection on the mutation rate itself. Indeed, the mutation rate is not only the input to heritable differences, but itself a phenotype, subject to genetic drift and selection (9) (10).

The mutation rate may also evolve as a byproduct of changes in the life history of the species, e.g., the ages at which males and females typically reach puberty and reproduce. Because the life history of the species modulates the length of exposure of the germ cell lineage to distinct developmental stages, shifts in life history can lead to evolution in the per generation mutation rate and spectra. A variant of this model has long been invoked to explain the observation that shorter-lived species (such as rodents) tend to have higher rates of neutral substitutions per unit time than longer lived ones (such as primates). The “generation time effect” posits that species that are shorter-lived accrue more mutations per unit time because they undergo more cell divisions per unit time (11–15).

Evaluating these possible explanations requires comparative data on germline mutation from closely related species that share much of their developmental program but differ in some key features. In principle, such data are now straightforward to collect, by resequencing genomes from tissue samples (for practical reasons, often blood) of mother, father and child and estimating the number of DNMs that occurred in that generation. This approach has been applied to human pedigrees in numerous studies, confirming that most mutations are paternal in origin and revealing a strong, linear paternal age effect as well as a weaker maternal age effect on the number of DNMs (16–18). Analogous data have also been collected in much smaller numbers of trios from other mammals, including mice (4), wolves (19), cattle (2), and primates (see Table 1 for primates).

Given the wealth of information for humans, comparisons to other primates may be particularly informative: this relatively closely related group of species varies markedly in their life histories, metabolic rates and other potentially salient factors, and there exist well-documented differences in their per year substitution rates (20,21). Among the handful of apes and Old World Monkeys (OWMs) from which direct estimates have been obtained, the estimated per generation mutation rate varies four-fold. This inter-species variation has largely been interpreted in light of differences in life histories and cell division rates among species [e.g., (22–24)].

These interpretations rely on the assumption that mutations track cell divisions, as expected in various settings, most obviously if mutations are due to errors in replication. The number of mutations then increases with each DNA replication cycle, at a rate that depends on the fidelity of DNA replication in that cell type (15). If instead mutations arise from damage that is inefficiently repaired relative to the length of the cell cycle, and the damage rate remains fairly constant, then the accumulation of mutations may track time (age) rather than numbers of cell divisions (15). In this case, differences among species may be more reflective of the damage rates that they experience and their typical ages at reproduction. While the relative importance of different sources of mutagenesis remains unknown, in humans, three lines of evidence suggest that many germline mutations are not tracking cell divisions: (1) there is a maternal age effect on mutation, which likely reflects (at least in part) damage accumulating in oocytes (16,17); (2) the mutation spectrum of mutations in males and females is similar (16,25,26), which one might not expect if the sources of mutation in the two sexes were largely non-overlapping; and (3) the ratio of male to female mutations barely increases with parental ages, despite the fact that oocytogenesis is completed long before reproduction whereas spermatogenesis is ongoing throughout life (27). Given these observations from humans, it is unclear what one should expect when comparing them to other species.

Here, we aimed to compare mutation patterns in humans to those in olive baboons, an Old World Monkey with a younger reproductive age. This comparison is of interest because male baboons typically enter puberty (and hence spermatogenesis) around age 6 and reproduce four years later, with an estimated 33 spermatogonial stem cell (SSC) divisions per year, whereas human males enter puberty by age 13 and reproduce on average 19 years later, with an estimated 23 SSC divisions per year (28–33). While there is considerable uncertainty about these numbers, they suggest that human sperm may be the product of 2.4-fold more cell divisions post-puberty relative to baboon sperm. This inter-species contrast thus sets up two predictions: if mutations are replicative in origin and rates per cell division have not evolved, we expect fewer mutations to arise from spermatogenesis in baboons compared to humans; in addition, there should be a stronger paternal age effect in baboons.

A number of technical difficulties stand in the way of this comparison. Germline mutations are rare (on the order of 100 in a 6 Gb genome), and reliable estimates are therefore exquisitely sensitive to false negative and positive rates, which in turn depend on sequencing coverage, the species’ reference genome quality, and the variant calling pipeline. Within humans, different studies have arrived at discrepant estimates of parental age effects, mutation spectra, and mutation rates at a given paternal age (34); indeed, significant differences are seen even between estimates based on two-versus three-generation families within a single study (27). These observations strongly suggest that comparisons among species are likely to be confounded by the choice of analysis pipeline. To minimize these technical biases, we resequenced three-generation families, calling putative DNMs in the second generation and then verifying transmission and phasing mutations with the third; moreover, we applied the same variant calling pipeline to the two species in orthologous regions of the genome.

## Results

### Identifying DNMs from Pedigrees in Parallel in the Two Species

In total, we resequenced the genomes of 26 humans from three three-generation pedigrees and 29 olive baboons from two three-generation pedigrees, to mean depths of 24X and 45X, respectively (Fig. 1). After mapping the reads of each individual to its species’ reference genome, we called variants using GATK following standard quality control practices (35). We identified single nucleotide polymorphisms (SNPs) at which the F_1_ child was heterozygous whereas both P_0_ parents were called homozygous for the reference allele, thus constituting a Mendelian violation. We filtered these variants on a variety of quality statistics and sequence metrics (see Methods) using the same criteria in both species, thereby identifying 1,046 putative autosomal DNMs in the 10 focal human F1 individuals and 497 DNMs in the 12 focal baboon F1 individuals (Fig. 1). We additionally flagged putative DNMs occurring in clusters, which we defined as DNMs arising fewer than 100 bp away from a neighboring DNM. Analyzing transmission patterns and validation results from Sanger sequencing, we concluded that clusters containing more than two variants were likely artifacts and thus discarded them in subsequent analyses (see Methods, Fig. S1). After this filtering step, we retained a total of 971 human and 465 baboon putative DNMs.

**Figure 1.**
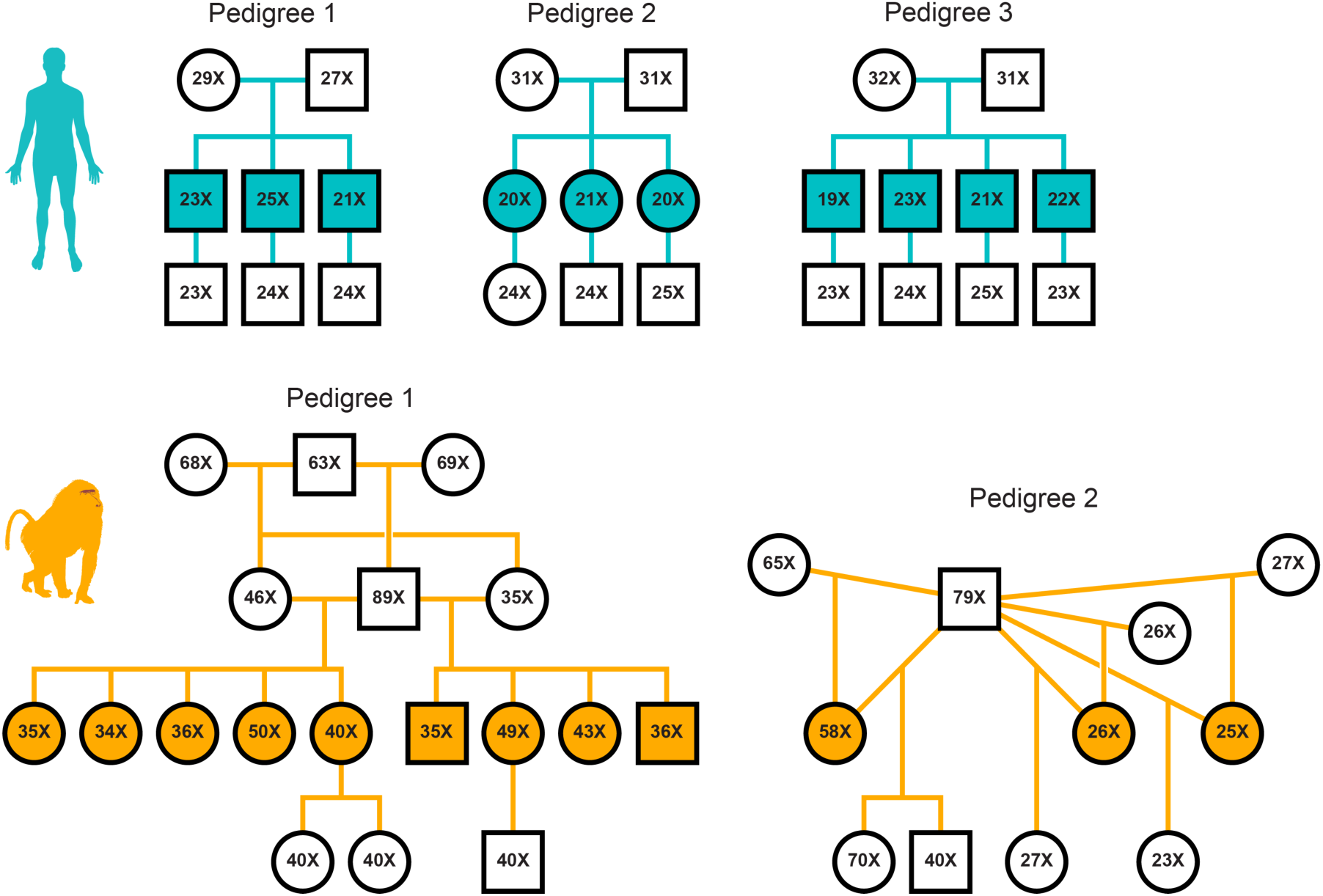
Pedigrees and mean depth of coverage of the sequenced individuals. Relationships among individuals in three human families (top row, teal) and two extended baboon pedigrees (bottom row, orange) are shown. Squares and circles represent males and females, respectively, and are annotated for the mean depth of sequencing coverage of the individual. Filled shapes indicate the focal F_1_ individuals in which mutations were called. Sample identifiers and birth dates for all individuals are specified in Table S2–S3.

By applying the same filtering pipeline to both species, we aimed to create a reliable comparison of their mutational properties. Furthermore, we identified regions of the genome that are orthologous between humans and baboons and in which we were highly powered to detect DNMs in all individuals, based on sequencing depth (see Methods). After restricting our analysis to these orthologous regions, which spanned a combined total of ~1.6 Gb, 503 putative human DNMs and 249 baboon DNMs remained. Rates in the orthologous regions are only slightly lower than those estimated using the entire callable genome (for which the mean across trios is ~2.5 Gb in human and ~2.3 Gb in baboon; see below).

Even when relatively infrequent, spurious DNMs can have a non-negligible impact on the mutation spectrum and the estimated fraction of DNMs that are of paternal as opposed to maternal origin (bringing it closer to 1:1 than it is truly). We therefore ran a series of analyses to estimate false negative rates (FNR) and false discovery rates (FDR). First, we generated a set of 20,000 simulated DNMs in each trio in such a way as to create Mendelian violations that mimic de novo mutations, then ran this set of simulated DNMs through our filtering pipeline and assessed the rate at which we failed to detect the mutation in the proband. This approach yielded median FNRs across trios of 9.80% (range: 8.96–17.95%) in humans and 7.93% (range: 7.05–9.51%) in baboons (Fig. S2A).

To measure trio-specific FDRs (i.e., the proportion of spurious mutations in our putative DNM call sets), we exploited the fact that false positive DNMs identified in an F_1_ individual are very unlikely to be observed in the F_2_ offspring, whereas genuine DNMs should be observed approximately half the time. In practice, we calculated FDRs using a maximum likelihood based model (see Methods), which accounts for the status of DNMs in the F_2_ generation (i.e., observed or not) as well as the probability of observing transmission events given our approach to calling variants and F_2_ coverage. We estimated median FDRs of 20% in humans and 18% in baboons (Fig. S2B). In parallel, we Sanger-sequenced putative DNMs called in one human individual both inside and outside of the orthologous regions in humans and baboons (Table S1, Fig. S3). Our transmission-based approach yielded an estimated FDR of 0% (95% confidence interval [CI]: 0–21%) for this individual. Consistent with that estimate (p-value = 0.35, by a Fisher’s Exact Test), we found only one false positive out of the 24 DNMs that we were able to check by Sanger sequencing (Fig. S4). In what follows, we used the transmission-based FDR for each trio.

### Estimating Sex-Specific Germline Mutation Rates and Age Effects

In order to assess maternal and paternal effects on mutation, we determined the parent-of-origin of the putative DNMs using a combination of read tracing and pedigree-based phasing. In humans, we were able to phase a total of 182 DNMs by the first approach and 220 by the second, with highly concordant parental assignments for mutations that could be phased by both (76 of the 77 DNMs). We then combined the phasing information with our estimated FDR and FNR values to infer the total number of maternally or paternally derived DNMs for each F_1_ (scaled up to an autosomal haploid genome size of 2.881 Gb, based on the hs37d5). Performing a Poisson regression for each sex separately of human DNM counts against parental ages at conception, we found a significant (p-value < 0.001) paternal age effect of 1.00 (95% CI: 0.36–1.63), as well as a significant (p-value < 0.001) maternal age effect of 0.79 (95% CI: 0.47–1.11).

Our estimate of the paternal slope, which is based on our sample of only 10 human trios (Fig. 2A), is consistent with the one reported by Gao et al., who relied on a sample of 225 three-generation trios (Likelihood Ratio [LR] test p-value = 0.258; matching the genome size surveyed) (16,27). Our maternal slope, however, is slightly higher (LR test p-value = 0.022). A possible explanation is that our sample includes two trios with mothers over 40. Indeed, Gao et al. report a better fit of an exponential than linear model, driven by observations for mothers over 40, and likewise such a model fits our data slightly better than their linear model (ΔAIC [Akaike information criterion] = - 2.94). In turn, our sex-averaged germline mutation rate is 1.25×10^−8^ (95% CI: 1.15– 1.34×10^−8^) per bp, in agreement with previous estimates of the human mutation rate (16,25,27,36,37), for paternal ages of 32.0 years and maternal ages of 28.2 years, the mean generation time in a previous DNM study that sampled over 1,500 trios (16). Together, these comparisons to larger data sets in humans suggest that our DNM filtering and discovery pipeline is reliable.

**Figure 2.**
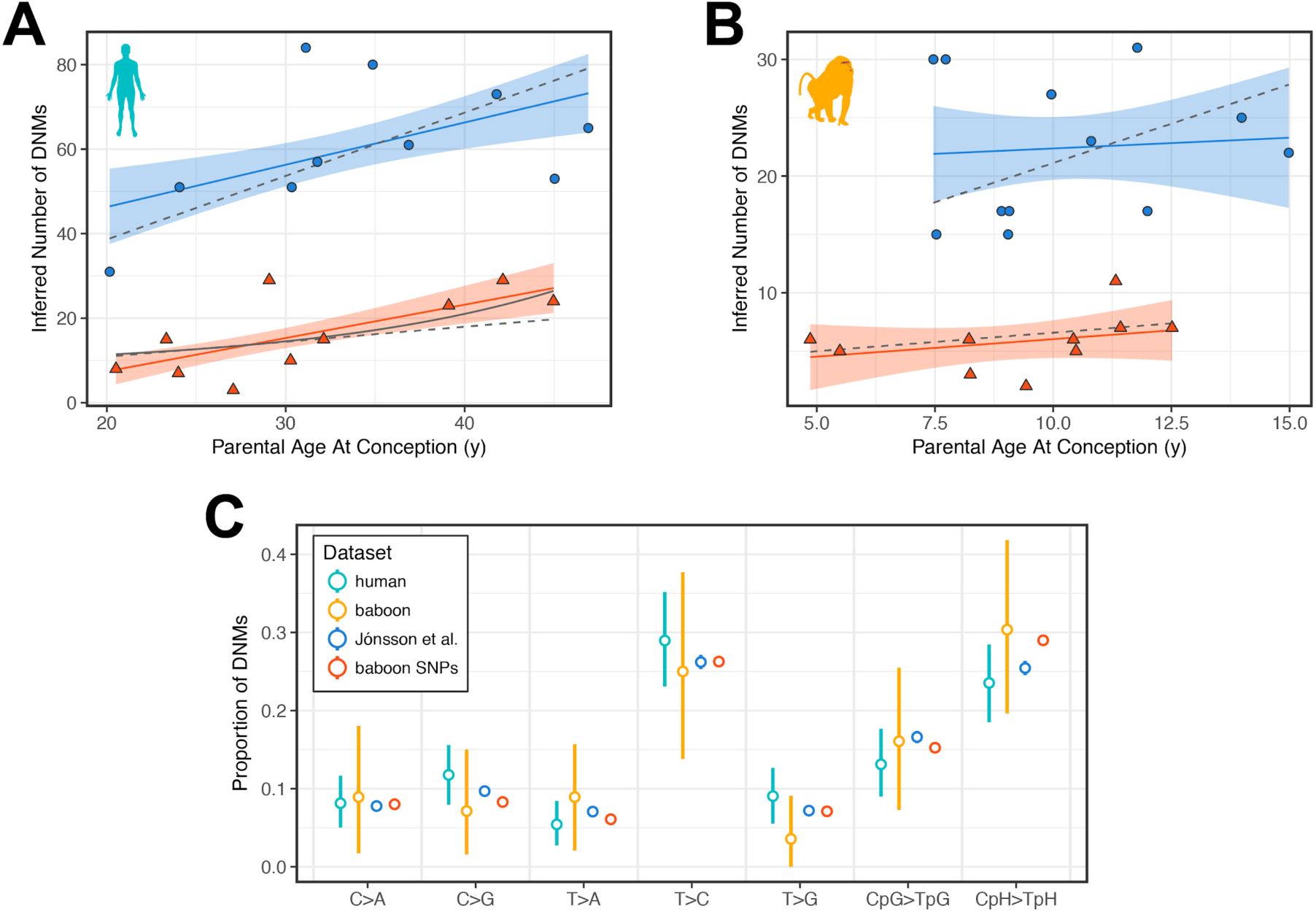
Dependence of mutation count on sex and age. Each human (in A) and baboon (in B) F_1_ individual is represented by a blue circle and a red triangle, which indicate how many DNMs were estimated to have occurred on the paternal or maternal genomes, respectively. Maternal points for two baboons were omitted due to lack of information about the age of the mothers. Solid blue and red lines indicate the best-fit obtained from a Poisson regression, for each sex respectively. Blue and red shaded regions denote the 95% confidence intervals associated with the regression coefficients. Grey dashed lines represent the sex-specific fits from Gao et al., (27), scaled to account for differences in genome size in (27) versus this study. In A, the solid grey line denotes the exponential maternal age effect fit from Gao et al. We did not find a significant difference between the parental slopes of the two species (LRT p-value = 0.22 in fathers, 0.12 in mothers); we therefore used the human estimates of age effects for baboons, as they are much more precise (27). (C) Human and baboon mutation spectra, in our sequenced cohorts and previously published datasets. Spectra are shown for: all transmitted DNMs in humans (teal); all transmitted baboon DNMs (orange); DNMs called in a large set of Icelandic three-generation pedigrees (16) (blue); and low-frequency SNPs (non-singleton alleles at frequency < 5%) identified in a sample of distantly related baboons (red). Each point denotes the relative proportion of mutations that belong to one of seven types of mutations, as indicated on the x-axis; reverse complement mutation types were collapsed together into a single type (e.g., the left-most column, C>A, includes G>T mutations). Transitions at cytosine sites were separated into those that occurred at a CpG (CpG>TpG) or a non-CpG (CpH>TpH) site. For all datasets, only DNMs and SNPs located within regions of the genome determined to be orthologous between humans and baboons were included. Vertical lines represent 95% confidence intervals from bootstrap resampling of 50 cM blocks in humans and 50 Mb blocks in baboons. Confidence intervals on the two reference datasets (blue and red) were constructed by bootstrap resampling of mutations but are small and hidden by the points.

We therefore proceeded to estimate the same parameters in our set of baboon DNMs by the same approach. As before, we inferred sex-specific DNM counts in each F_1_ by two phasing approaches and adjusted counts by estimated FDR and FNR. Given the higher diversity in baboons, we were able to phase a higher proportion of mutations by read-back phasing (144/249 = 58%) and (given the lower number of third generation individuals) we could phase only 56 by transmission; all but one of the 33 DNMs phased by both were in agreement. We then performed a Poisson regression against age at conception of the parents (Fig. 2B). Assuming that the mean ages of wild baboon fathers and mothers are 10.7 years and 10.2 years, respectively (Jenny Tung, personal communication), the per generation mutation rate in baboons is 0.55×10^−8^ (95% CI: 0.50–0.62×10^−8^) per bp, approximately 44% lower than the rate that we measured in humans. Reanalyzing the data considering all callable regions, not only orthologous ones, yielded a similar ratio between the two species, with a per generation mutation rate of 1.34×10^−8^ per bp in humans (95% CI: 1.25–1.44×10^−8^) and 0.59×10^−8^ (95% CI: 0.53–0.66×10^−8^) in baboons (Fig. S5).

We additionally compared the distribution of single base pair mutation types (i.e., the mutation “spectrum”) across our sets of human and baboon DNMs (Fig. 2C). We focused our analysis on the subset of DNMs that were observed at least once in the F_2_ generation (total of 221 in humans and 56 in baboons), reasoning that these were likely to be genuine DNMs. We classified each DNM into seven disjoint mutation classes: C>A, C>G, T>A, T>C, T>G, CpG>TpG, and CpH>TpH; the latter two classes denote mutation types occurring within and outside of a CpG context, respectively. In humans, the mutation spectrum in our trios was not significantly different from proportions obtained from a set of 14,629 human DNMs identified from 225 three-generation pedigrees by Jónsson et al. (16). Similarly, we did not observe any significant differences in mutation spectra in baboons when compared against low-frequency SNPs, which should closely reflect the de novo mutation spectrum. These results provide further evidence for the reliability of our calling pipeline. Although we lack power, we did not identify a significant inter-species difference in proportions for any of the mutation types (unadjusted *χ*^2^ test p-value > 0.05 for all comparisons).

Our findings in baboons thus suggest that the per generation mutation rate in OWMs is approximately half that of great apes, in which direct estimates range from 1.13×10^−8^ per bp in gorillas to 1.68×10^−8^ per bp in orangutans (23). Existing point estimates for OWM species are generally lower than those of apes, varying from a point estimate of 0.37×10^−8^ per bp in rhesus macaques to 0.94×10^−8^ per bp in green monkeys (24,38). While variation in mutation rates across primates is expected, these numbers are based on small numbers of trios and are thus highly imprecise (see Table 1). More critically, differences in how studies chose to filter DNMs and estimate error rates currently hinder reliable conclusions about how mutation rates differ across species. We sought to minimize these issues by identifying DNMs in a parallel manner across two species in the same regions of the genome, thus strengthening the case for a substantially lower mutation rate per generation in OWMs compared to apes.

### The Strength of Male Germline Mutation Bias is Similar in Baboons and Humans

We considered sex-specific mutation rates in the two species, comparing observations to what we would predict under an extreme model in which all germline mutations are replicative in origin. We further assume that human and baboon ova are the product of the same numbers of cell divisions (and the same per cell division mutation rates) and that SSC division rates are 23 cell divisions per year estimated in humans versus 33 in baboons (29,32). In females, we then expect similar per generation mutation rates in the two species; instead, the mutation rate is 0.24×10^−8^ (95% CI: 0.20–0.28×10^−8^) at age 28.2 in humans and 0.12×10^−8^ (95% CI: 0.088–0.15×10^−8^) at age 10.2 in baboons, a two-fold difference. Given recent evidence in humans that a substantial proportion of female mutations arise from damage to oocytes (16,18,27), the elevated rates in humans could result from the substantially older age of human oocytes compared to those of baboons.

In fathers, if the mutation rate per SSC cell division is the same in the two species, we expect a 1.5-fold stronger age effect in baboons compared to humans, due to higher division rates. Instead we find that the baboon data are highly unlikely under a paternal age effect of that magnitude or greater (LR test p-value = 2.5×10^−3^). As far as we can tell with limited data, paternal mutations are accruing no faster with age in baboons than humans (LR test p-value = 0.045).

These assumptions lead to a third prediction about the male bias in mutation. By a reproductive age of 32 years, the human male germ cell should be the product of 474 cell divisions (assuming that no cell divisions occur between birth and puberty) as compared to the average of 31 cell divisions in the female germline; in contrast, a baboon male germ cell should results from 217 cell divisions, due to a much younger average age of reproduction and despite the slightly higher rate of spermatogonial cell division. The ratio of male to female mutations should thus be roughly 2.2-fold larger in humans than in baboons. To estimate the male to female mutation ratio, *α*, we again focused our analysis on the set of DNMs called in the F_1_ that were observed in the F_2_ generation. In our human samples, we estimated that 79.5% (95% CI: 73.8–84.6%) of the DNMs were paternal in origin—i.e., an *α* of approximately 3.9—which is consistent with multiple previous estimates for the same ratio of paternal to maternal ages as in our sample (16,18). In baboons, we obtained a paternal fraction of 76.4% (95% CI: 63.5–89.1%), or an *α* of 3.2, which is not significantly different from the value in humans (*χ*^2^ test p-value = 0.75). We obtained similar estimates for *α* by using point estimates from our regression fits at the average ages of conception described above (Fig. 3A). Moreover, we can reject a model in which *α* in humans is 2.2-fold greater than in humans (LR test p-value = 1.4×10^−6^). Thus, the degree of male bias in mutation is surprisingly similar between humans and baboons.

**Figure 3.**
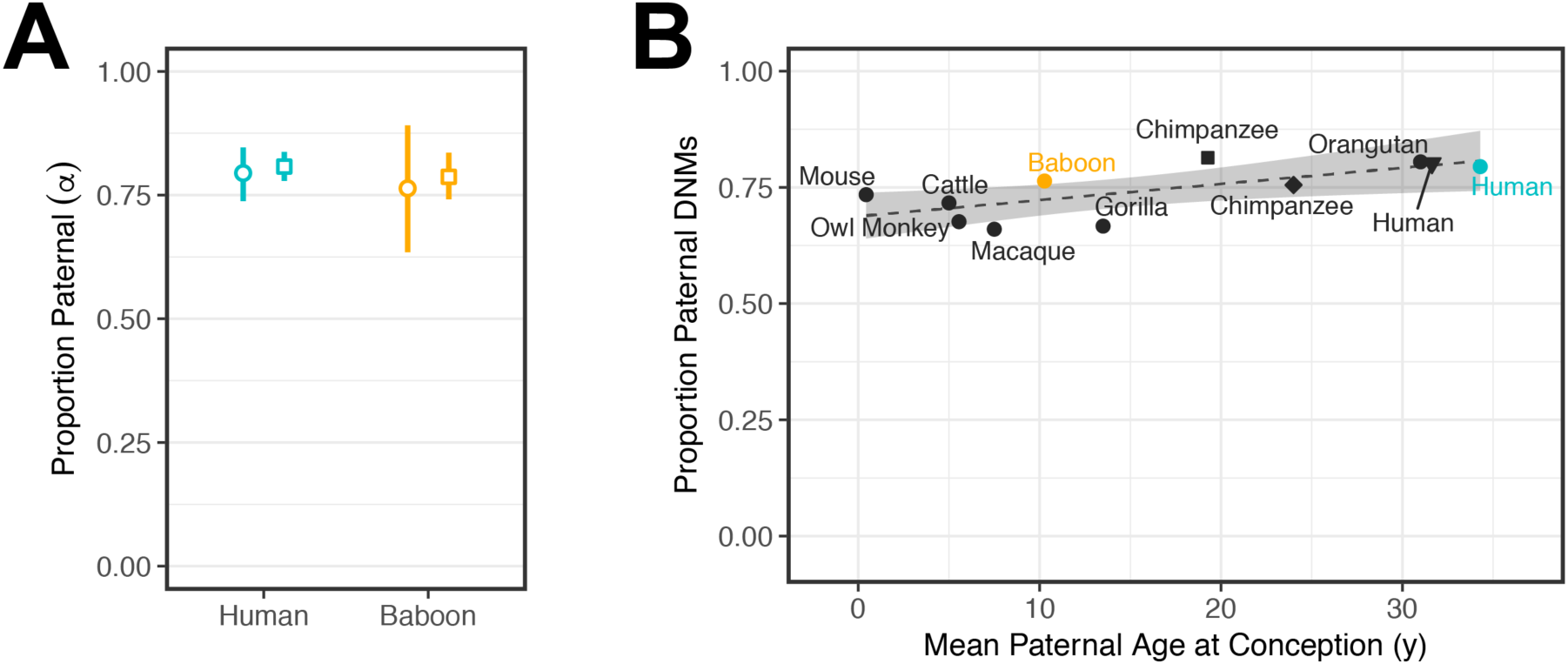
Estimates of the sex bias in mutation across mammals. (A) Ratio of paternal to maternal mutations, *α*, among phased DNMs. Circles indicate the fraction of DNMs assigned to the paternal genome among all transmitted DNMs (i.e., DNMs observed in an F_2_ offspring); in our samples, the mean ages in humans were 34.3 years for males and 31.3 years for females and in baboons, 10.3 years and 9.2 years, respectively (so the ratio of male to female generation time was about 1.1 in both). 95% confidence intervals were determined by bootstrap resampling of 50 cM blocks in humans and 50 Mb blocks in baboons. Squares denote the ratio of paternal to maternal mutations estimated for typical parental ages in the species, assuming ages 32.0 and 28.2 years for human males and females, respectively, and 10.7 and 10.2 years for baboon males and females. Vertical lines denote the 95% confidence intervals of the age effect and intercept estimates, obtained assuming normally distributed errors. (B) The ratio of paternal to maternal DNMs estimated from pedigrees across nine mammalian species as a function of the mean male generation times in the study (for references, see Table 1). The square and diamond points denote separate chimpanzee estimates from Besenbacher et al. and Tatsumoto et al., respectively (23,47). The black triangle denotes the estimate for DNMs phased by transmission in 225 three-generation human pedigrees from Jónsson et al. (16). Human and baboon estimates reported in this study are colored teal and orange, respectively. Fitting a linear regression (dashed line) yields an estimated slope of only 3.5×10^−3^ (95% CI: 1.0×10^−4^–6.8×10^−3^, p-value = 0.045). To avoid duplicates, values from one chimpanzee [Tatsumoto et al. (47)] and one human [Jónsson et al. (16)] study were not included in the regression. The shaded area denotes the 95% confidence interval of the slope and intercept.

To explore this observation further, we compared our estimates of *α* to what has been reported from direct pedigree analyses of other mammals, whose paternal generation times vary markedly, from months to years to decades. Although all these studies are based on scant data, the point estimates of *α* are remarkably similar in all seven species, with almost no increase with paternal age at reproduction (Fig. 3B). This observation echoes what is seen within humans, where despite the increase in the number of SSC divisions with paternal age, the male bias is already 3:1 by puberty and is relatively stable with parental ages (27).

## Discussion

### Implications for the genesis of mutation

We set out to test whether baboon and human mutation rates reflect differences in their life history, under a simple model in which mutations are proportional to cell divisions, females had the same number of cell divisions in the two species, and per cell division mutation rates over ontogenesis were the same in the two species. In this scenario, we would expect a stronger paternal age effect in baboons and a higher male mutation bias in humans. Neither of these expectations are met: the paternal age effect is not discernably stronger in baboons and the male bias is similar, as it is in other mammals for which direct pedigree estimates exist (Fig. 4).

**Figure 4.**
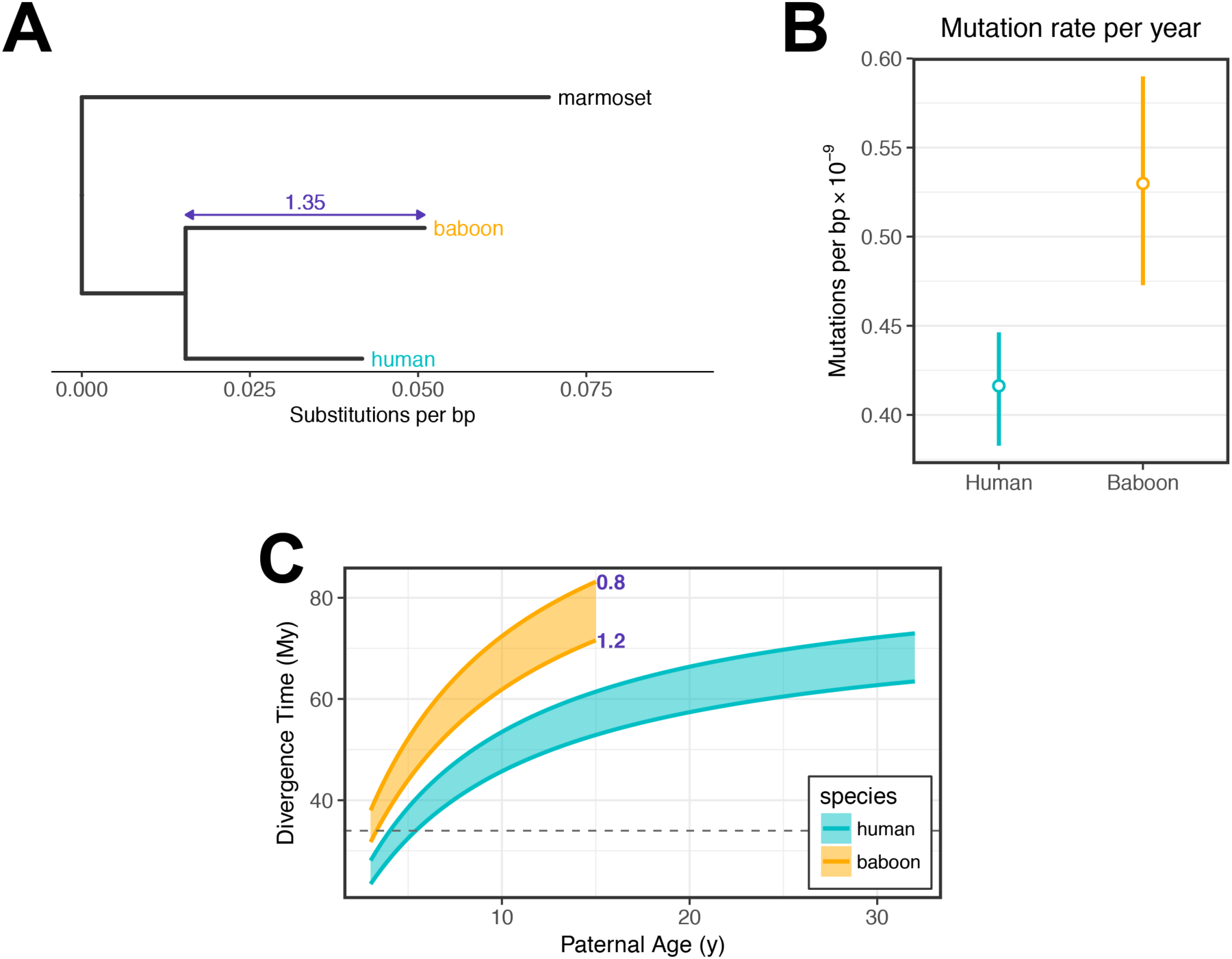
Implications of present-day human and baboon mutation rates for the evolution of the yearly mutation rate. (A) Phylogenetic relationship between humans and baboons with a marmoset (New World Monkey) outgroup. Branch lengths denote the substitution rate per base pair since the OWM-marmoset split as measured using data from (21) for all mutation types at putatively neutral regions of the genome. The relative branch length difference between baboon and human lineages is indicated in purple. (B) Sex-averaged mutation rates per year. Mutation rates were based on fitted values for typical generation times (i.e., assuming 32.0 and 28.2 years in human males and females, respectively, and 10.7 and 10.2 years in baboon males and females), and turned into a sex-averaged per year mutation rate following (36). Vertical lines indicate the span covered by the 95% confidence intervals of the intercept and slope of the age effect regressions. (C) Predicted divergence times of humans and OWMs as a function of parental ages. Divergence times were predicted using mutation and substitution rates measured in humans (teal) and baboons (orange), across a span of plausible past generation times. Each point within the shaded areas represents the divergence time calculated at a particular paternal generation time (x-axis) and paternal-to-maternal generation time ratio (ranging from 0.8 to 1.2) as indicated in purple. The dashed grey line indicates a plausible upper bound for the split time inferred from the fossil record (43,44).

A likely possibility is that one or more of the assumptions are wrong. In particular, a recent review argued that mutations are replicative in origin and hence track cell divisions but that the rate of SSC divisions is much lower than previously believed (39). While plausible (39,40), this explanation alone would not explain our findings: if rates of SSC divisions were very low, we would expect human and baboon mutation rates per generation to be highly similar when they are not.

Moreover, our findings add to a growing set of observations that do not readily fit a model where most germline mutations track cell divisions, including that: (1) in humans, there is a maternal age effect on mutations, which contributes a substantial proportion of maternal mutations, despite the absence of cell divisions after the birth of the future mother (16,18,27); (2) the male mutation bias in humans is already ~3:1 by puberty, when germ cells from the two sexes are thought to have experienced similar numbers of cell divisions by then (27); (3) the male mutation bias barely increases with parental age, and even less so once CpG transitions are excluded, despite ongoing SSC divisions (27); (4) OWM mutation rates are about half that of apes (this paper; 23) and (5) the sex bias in mutation rates is roughly similar across mammals (this paper; 14).

Observation 1 indicates that some fraction of mutations in females are non-replicative. Observations 2–5 could be explained by replication errors if we assume that a number of current beliefs about spermatogenesis are incorrect: namely that there are vastly more (or more mutagenic) cell divisions in males than females before puberty and that there are fewer (or less mutagenic) cell divisions in males after puberty. Even if both conditions hold, all the parameters would still need to conveniently cancel out both within humans and across species to generate an apparent dependence on absolute time rather than on cell divisions.

An alternative is that germline mutations are predominantly due to damage in both sexes. Accounting for all five observations would require damage to accrue at a relatively fixed rate and be inefficiently repaired relative to the cell cycle length (15), but at a somewhat higher rate in males than in females. Observation 4 additionally requires damage rates to be similar across mammals. This hypothesis would then explain the relative stability of the male bias in mutation in humans, the similarity of the male sex bias across mammals, and the similarity in the mutation spectrum in the two sexes. It would also explain why mutation rates per generation appear to reflect the typical ages of reproduction rather than numbers of cell divisions. While this hypothesis is parsimonious, it raises the question of why per year mutation rates vary across species and are higher in shorter-lived ones. One possibility is that rates of damage co-vary with life history traits (41). This explanation remains to be more explicitly modeled and evaluated in light of systematically collected comparative data from humans and other species.

### Comparing Contemporary Mutation Rates with Divergence Estimates

Studies of divergence in primates clearly demonstrate that neutral substitution rates vary substantially across the phylogeny (21). Notably, the olive baboon has accrued 35% more substitutions along its lineage as compared to humans since their common ancestor (Fig. 4A). If neutral, mutations are expected to fix at the rate at which they arise. Thus, we would expect the mutation rate per unit time in baboons to be substantially higher than that in humans. To evaluate this hypothesis, we converted our de novo germline mutation rates to yearly rates using a sex-averaged model that accounts for sex-specific relationships of mutation rates to age and sex-specific life history traits (36). This yielded yearly mutation rates of 5.30×10^−10^ in baboons, 27% (95% CI: 12–46%) larger than the rate of 4.16×^−10^ in humans (Fig. 4B). Thus, the ratio of present-day yearly mutation rates appears to be consistent (within error) with the observed substitution rate ratio in these two species.

Since biased gene conversion on mutation acts like selection and influences the substitution process, we further broke up substitutions by type, depending on whether the fixation probability was increased, decreased or unaffected by biased gene conversion; our findings are as expected, with mutations favored (or disfavored) by biased gene conversion showing slightly higher (or lower) substitution rates relative to mutation rates (Fig. S6).

If mutations are neutral and accumulate at a fixed rate, we can relate divergence levels to mutation rates in order to estimate the mean time to the most recent common ancestor (MRCA), in this case of OWMs and great apes. Dividing the human neutral substitution rate by the yearly mutation rate yields a time of 63 million years (My) while the same calculation using rates estimated in baboons yields a divergence time of 67 My. While an estimate of the mean time to the MRCA rather than of the split time, these two are expected to be very similar for species so diverged (42). Yet evidence from the fossil record dates the OWM-great ape population split time to at most 35 My (43,44).

A possible explanation for this discrepancy is that following the human-baboon split, yearly mutation rates in both species have slowed towards the present due to changes in life history traits (11,20,36). We explored this hypothesis by examining the effect of historical generation time on the inferred mean time to the common ancestor (Fig. 4C). We varied paternal generation times ranging from a lower bound of 3 years (the average age of reproduction of various New World Monkey species; 44) in both species to upper bounds of 32 years in humans and 15 years in baboons; we further allowed the male-to-female generation time ratio to vary from 0.8 to 1.2 (36). We used the age effect point estimates published by Gao et al. (27) as the age effect parameters of humans in our model, reasoning that—being drawn from a larger sample—these estimates would be more accurate in humans. We additionally used the Gao et al. estimates to model mutation accumulation in baboons, given that they were consistent with our data and that we had too few baboon trios to estimate parameters precisely (Fig. 2B). Our analysis suggested that, in both species, implausibly low generation times of approximately 3–5 years would be required to yield divergence times that are more in line with fossil-based estimates and even then, barely so. Thus, our results generalize the puzzle first pointed out by Scally and Durbin (46) of an apparent disconnect between the evolutionary times suggested by phylogenetic and pedigree data in humans. Reconciling them now requires not only a slowdown of the mutation rate per generation in humans but also in baboons.

## Data Access

Pedigree and DNM data are included in the Supplementary Tables and Data. Code for reproducing the figures will be published at https://github.com/flw88/baboon_dnms. Whole genome sequence data will be made available through the NCBI database of Genotypes and Phenotypes (dbGaP) and the NCBI Sequence Read Archive (SRA). Data for a subset of the sequenced baboons are currently accessible (NCBI BioProject accession number PRJNA433868).

## Acknowledgements

We thank Katherine Naughton for help preparing DNA samples for sequencing; Peter Andolfatto and the Andolfatto lab for help with the validation experiment; Ipsita Agarwal, Ziyue Gao, Guy Amster, and Arbel Harpak for comments on the manuscript; and members of the Przeworski and Sella labs for useful discussions. This work was supported by NIH grant R01GM122975 to M. P.; a Burroughs Wellcome Fund Career at the Scientific Interface award and Alfred P. Sloan Foundation Fellowship to P.M.; NIH grants R01HD21244 and R01HL085197 to C.O.; and NIH grant R24OD017859 to J.D.W. and Laura Cox. F.L.W. was supported in part by NIH training grant T32GM008224.

## Methods

### Human DNA Samples and Whole Genome Sequencing

We resequenced the genomes of 26 Hutterite individuals (51,52). The DNA samples were derived from either whole blood (25 individuals) or saliva (one individual). We performed 100 bp paired-end sequencing of the PCR free libraries, one per individual, to an average of 24X coverage on the Illumina HiSeq 2000 short-read sequencing platform at the University of Chicago. All individuals consented to this use of their DNA sequences, which was approved by the Institutional Review Boards of the University of Chicago and Columbia University.

Our cohort consisted of three multi-sibling families, containing 10 distinct parent-child trios (Fig. 1). Throughout, the first generation is referred to as P_0_, and the focal generation F_1_. We also sequenced one offspring of each of the F_1_ individuals, referred to as F_2_, which allowed us to assess transmission of DNMs to a third generation. Ages of both P_0_ parents at the birth of the F_1_ offspring were recorded for all 10 trios (see Table S2).

### Baboon DNA Samples and Whole Genome Sequencing

Our samples of olive baboon (*Papio anubis*) individuals originated from the captive colony housed at The Southwest National Primate Research Center (SNPRC), which was established in the 1960s with founder animals from southern Kenya. We initially sequenced a single sire, two dams, and their nine F_1_ offspring (Pedigree 1 in Fig. 1), obtaining baboon DNA samples extracted from liver or buffy coat from the SNPRC. A paired-end 150 bp library was prepared for each sample using the TruSeq DNA PCR-Free Library Prep Kit (Illumina), and sequenced on the Illumina HiSeq X platform. We expanded our initial two-generation cohort by including sequences generated by a parallel project at the SNPRC on the Illumina HiSeq 2500 platform (53). Thus, we analyzed a total of 26 baboons distributed into two pedigrees. The P_0_ generation is comprised of three dams and two sires that have 12 F_1_ offspring between them. F_2_ offspring are also present for a subset of these F_1_ baboon individuals. Pedigree 1 also includes the parents of the P_0_ individuals. We obtained an average coverage of 39X in the F_1_ generation and 52X in the P_0_ generation (Supplementary Data).

### Calling SNPs and Indels

We mapped our human (Hutterite) and baboon DNA sequences to the hs37d5 and papAnu2 reference genomes, respectively. To do so, we first interleaved the FASTQ formatted paired end reads with seqtk mergepe (version 1.2-r101-dirty) and trimmed sequencing adapters with trimadap (version r11) (54,55). Having thus prepared the sequences, we next used the Burrows Wheeler Aligner, bwa mem (version 0.7.9a-r786), to align the human and baboon reads to the hs37d5 and papAnu2 reference genomes, respectively, and added read groups as needed (56). We further processed the aligned sequences by marking duplicates using samblaster (version 0.1.24) and then sorting the alignments with samtools sort (version 1.3) (57,58). Finally, we used MergeSamFiles and MarkDuplicates from Picard Tools (version 2.9.0) to merge read groups by sample and perform a final round of duplicate marking, respectively, resulting in a BAM file of processed alignments for each individual. Default parameters were used for all software described above, unless otherwise noted.

We further improved our read mappings by running them through GATK (version 3.7) IndelRealigner and Base Quality Score Recalibration, in keeping with the GATK Best Practices workflow (35,59). These two steps benefit from the inclusion of sets of confident SNPs and/or indels. For humans, we used dbSNP build 138 available from the GATK resource bundle for the calibration. Genomic resources are more limited for *P. anubis*, and we therefore did not have access to a similar “gold standard” set of SNP and indel variation in baboons. To overcome this deficit, we generated two datasets to aid in calibration. We constructed the first dataset by making SNP and indel calls in our initial set of aligned reads using GATK’s HaplotypeCaller and GenotypeGVCFs tools. (We ran HaplotypeCaller with the pcr_indel_model parameter set to NONE to prevent the algorithm from incorporating PCR error-corrections on the variant likelihoods, since we sequenced our reads using a PCR-free protocol.) We then used bcftools isec (version1.3) to identify variants that were fully transmitted across all four generations of our baboon pedigree (i.e., variants in individuals 32043, 32849, and 33863 that were also observed in their respective parent, grandparents, and great-grandparents), reasoning that these are likely real polymorphisms (60). Our second dataset consisted of SNP and indel variants called from a sequencing panel of 89 unrelated baboons (49; available at NCBI BioProject accession number PRJNA433868), which we subjected to a number of site-level “hard filters” as suggested by the GATK Best Practices workflow (“QualByDepth” >2, “FisherStrand” < 60, “RMSMappingQuality” > 40, “MappingQualityRankSumTest” > −12.5, “ReadPosRankSumTest” > −8, “StrandOddsRatio” < 3 for SNPs) (35,53). The above procedure thus yielded one set of “known” variants for humans (from dbSNP) and two sets for baboons (from transmission and an outside sequencing panel).

For each species, we supplied the appropriate set(s) of known variants along with the initial set of mapped reads to GATK’s IndelRealigner and Base Quality Score Recalibration tools to produce reliable alignments for each individual. These alignments were in turn used to call SNP and indel variants using GATK HaplotypeCaller with the pcr_indel_model parameter set to NONE, as above. Finally, genotype likelihoods were jointly assigned across all 29 samples using GenotypeGVCFs (61).

### Identifying De Novo Mutations

Having called variants in both humans and baboons, we next focused on identifying de novo mutations (DNMs) in the trios available in our two cohorts. Specifically, we searched for point mutations present in the offspring (heterozygous for the ALT allele) that were absent in both parents (homozygous for the REF allele), thus constituting a Mendelian violation. Using bcftools query, we identified 41,164 such sites across our 10 human trios and 85,283 sites across our 12 baboon trios. As most of these sites are likely false positives, we identified possible sources of errors and developed a number of filters to increase our specificity.

First, we applied a number of site-level filters to avoid poorly sequenced or misaligned regions of the genome. For each individual, we constructed upper and lower cutoffs on the depth of read coverage by performing a two-sided Poisson test on depth at each site under the null hypothesis that the lambda (!) parameter was equal to the mean read depth of that individual (as reported by the GATK DepthOfCoverage tool). We required that, for a given trio, a site must have a read depth that yields a p-value > 2×10^−4^ in all three members of the trio to be considered callable. To this end, we identified the depth cutoffs of the rejection region and used these values as the minDepth and maxDepth arguments to GATK’s CallableLoci tool, which outputs bed files delineating callable regions. We used the bedtools intersect tool to intersect these regions across the members of each trio(62). In addition to filtering on depth, we required called SNPs to be biallelic and to have a quality (QUAL) score greater than 100. We also hard-filtered sites using the settings recommended by the GATK Best Practices workflow (“QualByDepth” > 2, “FisherStrand” < 60, “RMSMappingQuality” > 40, “MappingQualityRankSumTest” > −12.5, “ReadPosRankSumTest” > −8, “StrandOddsRatio” < 3 for SNPs) (35). We further removed known variants, filtering out SNPs if they were observed as variants in a panel of 87 Hutterite individuals (which did not include members of our sample) or the 89 baboon sample described above (53,63). For a given trio, we also filtered out SNPs if the same ALT allele was observed in an unrelated family.

We reasoned that false positive DNMs might be introduced by miscalling truly homozygous sites as heterozygous in the F_1_. To guard against this possibility, we required SNPs of a given offspring to have a genotype quality (GQ) greater than 40 and that the number of reads supporting the alternate (mutant) allele—i.e., the allelic depth (AD)—be greater than 3. We also performed, for each SNP, a two-sided binomial test on allelic balance (AB), the proportion of reads supporting the alternate allele, under the null hypothesis that the frequency = 0.5, and discarded SNPs with p-values less than 0.05. We note that a frequency of 0.5 implicitly assumes that the mutation is constitutive in all cells of the offspring rather than mosaic.

False positive DNMs might also be introduced due to miscalling truly heterozygous sites as homozygous in the P_0_. To remove such errors, for each trio, we required AD = 0 in at least one parent and GQ > 40 in both parents. We took the added step of removing DNMs determined to be in clusters, which we defined as groups of DNMs of size 3 or larger (see section below for our reasoning and details on cluster detection).

A comprehensive list of all filters listed above is available in Table S4. After filtering we were left with 971 and 465 DNMs in the human and baboon cohorts, respectively. For trios where a third generation individual was also sequenced, we determined DNMs to have been “transmitted” if at least 2 reads supporting the DNM were present (AD ≥ 2) in that individual.

### Estimating Size of the Accessible, Orthologous Genome

Obtaining a mutation rate per base pair requires calculation of the total number of base pairs for which our approach provides high power to call mutations, the so-called accessible genome size. To this end, we defined the size of an individual’s accessible haploid genome as the number of sites that were annotated as “CALLABLE” by the GATK CallableLoci tool and passed the known variation filters, as described above. For a given trio, we further restricted the accessible haploid genome to the intersection of accessible regions across all three members of the trio and removed sites if at least one parent had a non-homozygous reference genotype, resulting in a single bed file of accessible genomic regions for each trio. This procedure of subtracting and intersecting genomic regions was performed using the bedtools toolkit (62). (The size of the accessible diploid genome is then simply twice that of the haploid genome.)

In order to make interspecies comparisons in later analyses, we further identified the segments of the baboon and human genomes that are not only accessible but also orthologous. To do so, we first identified orthologous genomic regions using the Enredo-Pecan-Ortheus (EPO) seven-primate multi-alignment dataset (ENSEMBL release 76) (64,65). We used tools from the mafTools package to preprocess the original MAF-formatted files by: (1) removing duplicates using mafDuplicateFilter; (2) extracting human-baboon autosome multi-alignments using mafFilter; (3) coercing the multi-alignment block coordinates to use the positive (sense) Homo sapiens strand using mafStrander (66). We then used a custom awk script to read the processed MAF files and create a set of bed files containing the human and baboon reference coordinates of each pair of orthologous regions. Separately, we intersected the accessible regions across all trios in each species cohort, resulting in an accessible haploid genome for each species. We then used the liftOver tool to convert the accessible human coordinates from the original hs37d5 reference to GRCh38, the reference genome used in the EPO multi-alignment (67). Lastly, we performed a final round of intersections between each of the cohort-specific accessible genomes and our set of orthologous regions, resulting in a set of bed files defining a set of regions that were orthologous between the two species. The total size of the orthologous regions equaled 1,631,476,416 bp or about 64.6% and 69.7% of the mean callable genome of humans and baboons, respectively. Within the orthologous regions we identified 503 human and 249 baboon DNMs (equivalent to 51.8% and 53.5% of DNMs originally called across all callable sites).

### Performing False Negative Rate Simulations

Our stringent filtering process likely removed true DNMs, which would lead us to under-estimate the mutation rate. In order to measure the false negative rate (FNR) of our DNM-calling pipeline, we simulated a set of “real” DNMs and quantified the proportion of DNMs that were removed by our filters (Fig. S2A). Specifically, we created these simulated DNMs by searching our initial set of SNP variant calls for sites satisfying the following criteria: (1) the F_1_ and exactly one P_0_ parent (hereafter referred to as the heterozygous parent) of a given trio were both heterozygous for the alternate allele (genotype 0/1); (2) the other P_0_ parent was homozygous for the reference allele (genotype 0/0); (3) a P_0_ parent (hereafter referred to as the substitute parent) of the same sex as the heterozygous parent but from a different family was homozygous for the reference allele (genotype 0/0); (4) the site passed hard filtering and depth filters. Of the sites that met these criteria, we randomly sampled 10,000 sites. We then transformed these Mendelian transmissions into non-Mendelian DNMs by treating the substitute parent as the real parent of the offspring in question, in effect “replacing” the genotype and quality statistics of the true heterozygous parent with those of the homozygous substitute parent. Where multiple individuals could be used as the substitute parent, we selected one at random for each DNM.

In practice, we performed this substitution by removing the heterozygous parent as well as its descendants save for the focal F_1_ and F_2_s, and subsequently rerunning the joint genotyping algorithm (GATK GenotypeGVCFs) across the remaining individuals. Removing these related individuals and re-genotyping was necessary because transmitted heterozygous variants will be shared quite often with sibling F_1_s—much more often than is the case for true DNMs (2). Since the joint genotyping algorithm improves the genotype likelihood of an allele if it is observed in multiple individuals, the removal of other individuals that may carry the same allele mimics as closely as possible the status of a true DNM. The above procedure was performed twice for each trio, in one case replacing the mother and in the other case replacing the father, resulting in two sets of simulated DNMs for each trio.

Next we applied the remaining filters to the set of simulated DNMs, as if they were observed ones. The proportion of simulated DNMs that either failed the filters or were not retained during the re-genotyping stage served as our FNR estimate for each trio.

### Phasing De Novo Mutations

Having called our DNMs, we subsequently identified the parental backgrounds on which they arose by phasing the genotypes. To this end, we used both a read-backed phasing method, which takes advantage of the phasing information contained in short-read sequences, as well as a phase-by-transmission approach, which was feasible due to the three generations present in our pedigrees.

For each species, we performed read-backed phasing by feeding the appropriate SNP variant call-set VCF file into the GATK ReadBackedPhasing tool with the phaseQualityThresh parameter set to 20, producing a new VCF file in which phased SNPs were annotated with haplotype identifiers. We assigned a given haplotype to a particular P_0_ parental background if at least one SNP along the haplotype was unambiguously inherited from the parent in question. If more than one informative SNP was present in a haplotype, we required that all such SNPs indicate the same background; haplotypes that indicated conflicting parental backgrounds (i.e., if both maternally and paternally inherited SNPs were present) were given an ambiguous status. Using this procedure, we succeeded in identifying the parental backgrounds of 373 of the 971 human DNMs identified both within and outside of the orthologous regions. We successfully phased a higher proportion of baboon DNMs (266 of 465 total across the callable genome) as expected from their higher nucleotide diversity levels (68).

In order to phase DNMs by transmission, we employed a two-step process of pedigree-based phasing using the SHAPEIT2 (v2.r904) program then inferred the parental background from transmission observations in the third generation (69,70). We first converted our VCF-formatted SNP sets to BED/BIM/FAM format using PLINK-1.9 (71,72). We also filtered away SNPs if they failed our hard filters, had poor genotype quality (GQ < 20), were multiallelic, or had excess missingness across the set of samples (> 5%). Along with SNP observations, SHAPEIT requires a genetic map to weight the prior probabilities of transitioning between haplotype segments. For our human cohort, we used the HapMap Phase II b37 genetic map (73,74). Lacking a usable fine-scale genetic map for baboons, we opted to generate a uniform map assuming a rate of 2354 cM / 3100 Mb (75,76). We input our filtered SNP set and the species-appropriate genetic map into SHAPEIT, which we ran in the pedigree-aware duoHMM mode (flag -- duohmm), with a mean window size of 2 Mb (flag -W 2). This step generated both maximum likelihood haplotypes as well as a haplotype graph for each cohort. To obtain crossover event probabilities, we next sampled 10 haplotype sets from the haplotype graph by rerunning SHAPEIT with different seeds, using the standalone DuoHMM program to generate recombination maps for each sample, and averaging over the maps using mapavg.py (77). Finally, we determined the phase of any given transmitted DNM by identifying the backgrounds of nearby in-phase SNPs in the third generation that were unambiguously inherited from the proband; we assigned the transmitted DNM the opposite background if a recombination event with probability >50% was present before the first informative SNP on both sides of the DNM site.

For each DNM, we combined phasing calls from the read-backed and pedigree-based methods into a single consensus call (paternal or maternal). Trivially, if only one phasing method succeeded then we used that inferred haplotype; similarly, if both phasing methods assigned a DNM to the same haplotype then we chose that haplotype as the consensus phase. Across both orthologous and non-orthologous regions, we successfully phased a total of 143 DNMs in humans using both methods, of which 141 agreed in the haplotype call. We eliminated the two inconsistent DNMs from consideration when estimating *α* and inferring sex-specific DNM counts, under the assumption they are errors (see below).

In baboons, 51 DNMs were assigned haplotypes across the orthologous and non-orthologous genome using both the read-backed and pedigree-based techniques. Of these DNMs, three were transmitted to one F_2_ and disagreed in the assigned haplotypes; they were ignored in subsequent analyses. Baboon F_1_ individuals with two F_2_ offspring afforded two opportunities to infer phasing using transmission observations. We identified only one DNM (chr2:112,593,279 in F_1_ individual 7267) that was transmitted to two F_2_ offspring and pedigree-phased to both paternal and maternal haplotypes. To resolve this conflict, we manually assigned this DNM to the maternal haplotype, the phasing call indicated using the read-backed technique. Finally, five baboon DNMs exhibited “partial linkage” in that they were observed in exactly one of the two available F_2_ offspring despite those individuals having the same parental haplotype at the focal site. This pattern could arise if the mutation occurred during the development of the F_1_ (rather than being inherited from the P_0_ gametes), and thus it would not be identified every time its corresponding haplotype was observed (2). In agreement with this possibility, read-backed phasing was also available in three of the five DNMs and in each case the transmitted haplotype was inferred. Thus, we assigned these DNMs to the transmitted parental haplotype.

### Detecting DNM Clusters

For each trio, we defined a DNM as belonging to a “cluster” of DNMs if it occurred fewer than 100 bp away from a neighboring DNM in genomic coordinate space; a DNM cluster is then simply a set of two or more DNMs whose neighbor distances do not exceed 100 bp. Using this working definition, we identified 55 and 33 clusters in our human and baboon datasets, respectively (Fig. S1A). We considered a DNM cluster to be transmitted if we observed all the DNMs within the cluster transmitting to the F_2_ generation.

Across both species, we identified 22 clusters containing 3 or more DNMs in F_1_ individuals with an available F_2_ generation (26 clusters across all trios). If clusters behaved like singlet DNMs (i.e., having the same expected probability of transmitting in each trio; see “Estimating False Discovery Rates” below) then we would expect to observe ~12 of the 22 clusters in the F_2_ generation. In contrast, none of these clusters were observed to be transmitted (Fig. S1B; Fisher’s exact test p-value = 6.1×10^−5^). This finding agrees with the results from Sanger resequencing, as detailed below. In both species, DNMs co-occurring in clusters of size two (hereafter referred to as doublets) were transmitted to the F_2_ generation at a much lower rate than singlet DNMs. In particular, we observed a doublet in the H14 trio, which was both transmitted to its F_2_ child, H25, and verified as “present” by Sanger resequencing (we failed to verify any other DNM clusters in H14; see Fig. S4C).

Based on these observations from transmission analysis and Sanger verification, we reasoned that, within the context of our pipeline, clusters of different sizes harbor different false discovery rates (FDRs). As clusters of size 3 or larger appeared to have an extremely high FDR (consistent with 100%), we filtered these clusters out entirely from our DNM datasets, resulting in the removal of 75 human and 32 baboon DNMs. We subsequently proceeded to estimate FDRs separately for singlet and doublet DNMs in the two species.

### Estimating False Discovery Rates

We used the fraction of putative DNMs that transmit from F_1_ individuals to their F_2_ offspring to estimate, for each trio, the proportion of putative DNMs identified by our pipeline that are true positives—i.e., one minus the FDR. This approach relies on the fact that a spurious DNMs called in the F_1_ is exceedingly unlikely to be seen in the F_2_.

We first consider the case of singlet DNMs. Ignoring for now the fact that not all F_1_ baboons have F_2_ offspring, let *m_i_* ∈ {1, 2} be the number of F_2_ offspring present for the *i*^th^ F_1_. We assumed that each true positive DNM called in the *i*^th^ trio has probabilities *q_i,j_* of being observed in the *j*^th^ F_2_ (*j* ∈ {1, …, *m_i_*}) and that spurious DNMs will never be observed in the F_2_ generation. Treating the F_2_ status (observed or not observed) of each DNM as independent, the total number of DNMs observed in the *j*^th^ F_2_ is thus binomially distributed. If only a single F_2_ is present (*m_i_* = 1), then the number of observed singlet DNMs, 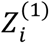, in the F_2_ is modeled as:

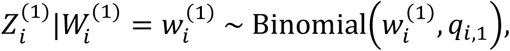

where the superscript (1) denotes singlet DNMs and 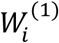 is the number of true positive singlet DNMs.

If two sibling F_2_s are present (*m_i_* = 2), then there are four outcomes (X) for the F_2_ generation: (A) the putative DNM can be not observed in both F_2_s, (B) observed in the first F_2_ but not the second, (C) observed in the second F_2_ but not the first, and (D) observed in both F_2_s. If we assume that the statuses of a DNM in two sibling F_2_s are independent, then the number of observed DNMs falling in each of the four categories can be modeled as:

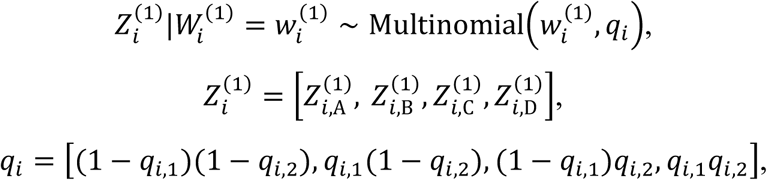

where 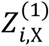 is the number of singlet DNMs observed for outcome X.

We made use of our simulated DNMs, which we described above in the context of estimating FNR, in order to obtain the probability *q_i,j_* of observing a DNM called in the *i*^th^ F_1_ in its *j*^th^ F_2_. We took this approach rather than assuming 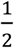 both because we do not have 100% power to detect a variant transmitted to the F_2_ (though in practice close to it for the accessible genome) and because GATK increases the likelihood weights of variants that are observed multiple times in a cohort, leading to deviation from the expected 50% probability of observing a called variant in the F_2_ at stringent filtering thresholds. Thus, for each F_1_-F_2_ pair, we estimated the proportion of simulated, filtered DNMs that were observed in the F_2_, denoted 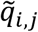.

Because the baboon pedigree includes loops (resulting from inbreeding between close relatives) for baboon trios with individual 1X2816 as the sire, we multiplied 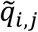 by a corrective factor of 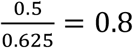 to account for the fact that simulated DNMs may be transmitted either through the F_1_ (as is typical) or through 1X2816 when it is the masked, heterozygous parent. The denominator of 0.625 is the sum of the joint probabilities of a randomly chosen SNP being transmitted and deriving from the P_0_ dam (0.25) or the P_0_ sire (0.375). Thus, formally:

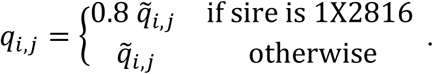

Using the parameterization outlined above, the likelihood of the observed singlet DNM data for trio *i* is thus:

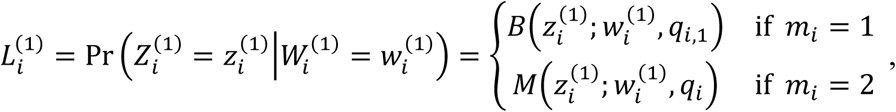

where *B* and *M* are the binomial and multinomial probability mass functions, respectively. Finally, we obtained a point estimate of the FDR of singlet DNMs for each trio using the maximum likelihood estimate of the number of true DNMs, 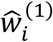:

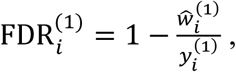

where 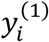 is the total number of singlet DNMs observed in trio *i*. In searching for the maximum likelihood estimate of 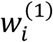, we constrain this quantity to be strictly less than or equal to the total number of singlet DNMs, 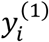, and greater than or equal to the total number of singlet DNMs observed in the F_2_ generation.

Seven of our baboon F_1_s lack any F_2_ offspring, which obviously precludes the use of transmission information for inferring a FDR for these individuals. For these individuals, we estimated an FDR by relying on the tight correlation we observed between the estimated FDR for individuals with an F2 and the mean coverage in F1s (Fig. S7). Specifically, we regressed the estimated FDR values on the F_1_ average depth of sequencing coverage; we then used the fitted linear regression to obtain point estimates of the FDR of baboon F_1_s without F_2_ offspring from their average depth of coverage.

For doublet DNMs, we calculated a single FDR across all trios, rather than a per trio rate, which was necessary given the rarity of doublets. We used the same framework described above for estimating FDR from transmission information for singlets, but aggregating variables over all trios in each species:

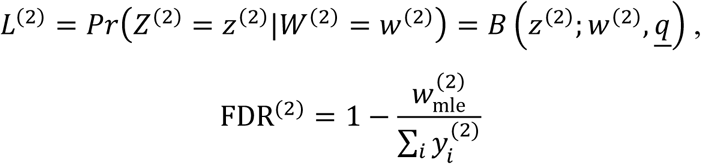

where *Z*^(2)^ is the number of doublet DNMs observed in the F_2_ generation, *W*^(2)^ is the number of true positive doublet DNMs, 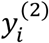 is the number of doublet DNMs observed in the *i*^th^ F_1_, and *<u>q</u>* is the arithmetic mean of *q_i,j_* taken over all F_1_-F_2_ pairs. All these operations were performed using the R statistical computing language.

### Sanger Sequencing

In order to assess whether our false discovery rates were reliable, we used Sanger sequencing to validate the presence or absence of DNMs called in one trio. Specifically, we designed primer pairs (Table S1) targeting all 89 DNMs called both inside and outside of the orthologous regions in H14 (Fig. S4). We also attempted to validate an additional 14 putative “cluster” DNMs in H14 that had been removed for being members of clusters sized three or larger. The clustering phenomenon allowed us to target multiple DNMs with a single primer pair such that we required only 87 primer pairs (designed using Primer3) to target the 103 total DNMs (78,79). We evaluated primer specificity using NCBI PrimerBlast (80). Using the high-fidelity Phusion polymerase (New England BioLabs E0553L), we attempted to PCR-amplify the 87 target sites in all three members of the H14 trio (H14, H11, H12). A subset of the primer pairs yielded nonspecific products, as indicated by multiple bands in gel electrophoresis. We performed band stab on these non-specific products, selecting the appropriately sized band and performing a second round of PCR with fewer cycles. In total, we succeeded in amplifying 54 sites. We performed a PCR product cleanup step to remove primers and small DNA products using Agencourt AMPure XP Beads (Beckman Coulter B37419AB) or spin columns (Qiagen 28104). Finally, we submitted the cleaned products as well as their respective forward primers to Genewiz for Sanger sequencing.

We visually inspected the resulting chromatograms using the SnapGene Viewer v4.3.5 and determined that Sanger resequencing succeeded in all three trio individuals for 25 of the 54 sites originally submitted (81). These 25 sites contained a total of seven cluster and 24 non-cluster DNMs. We verified that all seven cluster DNMs were absent in support of our decision to remove cluster DNMs from our dataset. In contrast, only one non-cluster DNM was verified as absent (see Fig. S4B–C for chromatogram examples), which at face value suggests a FDR of 4% (95% CI: 0–21%). We compared the distributions of validated versus spurious mutation counts in our Sanger results (23 versus one) against that inferred from transmission data. Of the 89 (non-cluster) DNMs observed in H14, 44 were observed in its offspring, H25, out of an expected 89*q*_H14,1_ = 89 × 0.492 ≈ 44, suggesting that there were no spurious DNMs. Comparing 1 out of 24 to 0 out of 44 by a Fisher’s exact test yielded a two-tailed p-value of 0.35 (Fig. S4A).

### Calculating Per Generation Mutation Rates and Age Effects

We inferred the number of “real” DNMs present in each trio offspring by adjusting the number of putative DNMs for false discovery rates (FDR), false negative rates (FNR), and the size of the accessible genome. The numbers of singlet and doublet mutations observed in the *i*^th^ F_1_ are 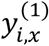 and 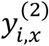, respectively, with the subscript *x* indicating the species. To obtain 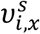, the error-adjusted number of mutations arising on a parental background of sex *x* in species (*s* ∈ {*female, male*}) in species *x* (*x* ∈ {*human, baboon*}), we used the relationship:

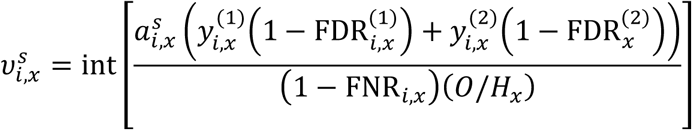

where 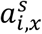 is the proportion of mutations in the trio on background of sex *s* 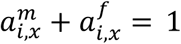 1); *0* is a constant denoting the amount of the genome that we determined to be orthologous and accessible (1,631,476,416 bp); *H_x_* is the haploid genome size of the species; and “int” is the nearest integer function. When an F_2_ is present, we estimated 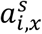 to be the proportion of transmitted mutations that were phased to background *s* (by both approaches or when no F2 was available, using read-back phased mutations only). As described above, we estimated 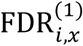 and 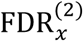 from F_2_ transmission observations and FNR*_i,x_* from DNM simulations. We obtained values for *H_x_* by counting autosomal base pairs in the reference genomes of each species yielding *H_hum_* = 2,881,033,286 and *H_bab_* = 2,581,196,250.

For each species, we model the number of de novo germline mutations inherited on a paternal (maternal) background as a function of the father’s (mother’s) age at conception, using a Poisson distribution:

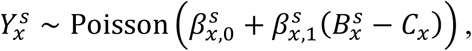

where 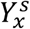 is the realized number of mutations; 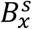 is the parental age at birth; *C_x_* is a constant denoting the typical gestation time of the species (Table S5); and 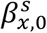 and 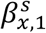 are the sex-specific intercepts and parental age effects. We fit this regression model separately for each sex and for each species using our error-adjusted DNM counts 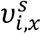 and the recorded parental ages at birth (Table S2–S3) of the P_0_ individuals, and obtain the maximum likelihood estimates 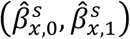 for the coefficients. Ages at birth were not available for baboon dams 1X0356 and 1X4519, and we therefore excluded their F_1_ offspring from the baboon maternal regression analysis. Regression computations were implemented in R using the glm function.

To predict the per base pair mutation rate at some average generation time 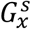 for each sex in each species, we considered:

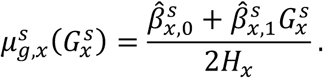

By summing maternal and paternal rates, we obtained species-specific mutation rates per base pair “per generation”:

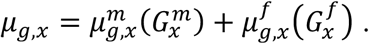

In the main text, we report mutation rate point estimates using paternal and maternal generation times thought to be typical of the two species.

For all mutation rates, we computed 95% confidence intervals by generating 1000 random draws of the regression coefficients 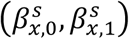 from normal distributions centered at the point estimates and taking the standard errors of the fit as the standard deviations. Each replicate draw provided point estimates for the mutation rate of interest, from which we generated a distribution.

### Timing of Life History Events in Baboons and Humans

We estimated the length of various life history stages of humans and baboons using data drawn from the literature. For both species, we obtained values for the typical gestation time (*C_x_*), male age at puberty, and male and female ages at reproduction 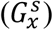, which are collated in Table S5. Wherever we were unable to find data for *Papio anubis*, we relied on estimates from other members of the *Papio* genus.

We drew typical gestation times from the AnAge database, using the value reported for *Papio hamadryas* (82). For typical male age at puberty in humans, we relied on an estimate of the median male age at spermarche by Nielsen et al. (30); in baboons, we based our estimate on the median age at testicular enlargement measured in a wild population of *Papio cynocephalus*, which maintains a hybrid zone with *Papio anubis* in the Amboseli Basin of Kenya (28). For typical male and female ages at reproduction, we used the mean paternal and maternal ages at conception of 1,548 Icelandic trios sampled in (16). For baboons, we used estimates derived from field data on parental ages at birth from the same wild Amboseli population noted above (Jenny Tung, personal communication).

### Quantifying Germline Cell Divisions

Estimating the number of germ cell divisions per generation is challenging, as it requires accurate knowledge of cell division rates across the multiple developmental stages that span from formation of the zygote to production of the mature gametes. The major stages of this process are currently understood to extend (1) from the first post-zygotic division to sex differentiation, (2) from sex differentiation to birth, (3) from birth to puberty, and (4) from puberty to reproduction (15,33).

In humans, a classic reference is Drost and Lee, who estimated that stage 1 is comprised of 16 cell divisions (33). They further estimated 21 cell divisions occurring during stage 2 in human males and 15 in females. Oocytogenesis is completed at the end of stage 2. In males, there are believed to be no cell divisions during stage 3. Under simplifying assumptions, the number of cell divisions in stage 4 depends on the length of the spermatogonial stem cell cycle (16 days in humans) as well as the typical age at puberty (approximately 13 years) (32). Assuming a typical male reproductive age of 32 years and taking into account the additional four cell divisions needed to complete spermatogenesis from spermatogonial stem cells yields 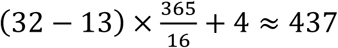 cell divisions for stage 4 in male humans. Assuming these numbers, the male germ cell is thus the product of approximately 474 cell divisions post-fertilization versus an average of 31 cell divisions in the female germline.

To the best of our knowledge, detailed analysis of germ cell division rates in baboons is limited to spermatogonial stem cells in stage 4, which are thought to divide every 11 days on average (29). Assuming typical male ages at reproduction and puberty of 10.7 and 5.41 years, respectively, there are then about 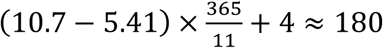 post-pubertal cell divisions in baboon males. For earlier developmental stages, we assumed that the number of cell divisions in stages 1–3 in the male baboon germline is the same as in humans, yielding a total of approximately 217 cell divisions. We discuss the impact of violations of these assumptions in the Discussion.

### Testing Age Effects and Intercepts

We tested for differences between the parental age effects measured in our human and baboon datasets by performing, for each sex, a Poisson regression on the combined data from both species. This regression model included a slope (age effect) and intercept term, as before, as well as an interaction term for the effect of species on the intercept. After fitting this three-parameter model with the glm function in R, we performed a likelihood ratio test to compare it to the four-parameter model previously obtained by regressing each species separately.

We also compared our estimated sex-specific mutation parameters to values obtained for the Jónsson et al. “deCODE” dataset by Gao et al. (16,27). To this end, we relied on estimates made using only the 225 deCODE trios for which a third generation was available, a similar study design to ours. This choice was motivated by the observation that mutation rate estimates based on three-generation pedigrees were different and likely somewhat more reliable than those based only on two generations (27). Since Jónsson et al. reported *R* = 2,682,890 bp as their callable genome, we multiplied the Gao et al. intercepts and age effects by a factor of *H_x_/R* to scale these values to the number of base pairs considered in this study (Supplementary Data). We then performed likelihood ratio tests comparing our fitted models to these adjusted Gao et al. parameter values in R. We note that the Gao et al. parameters were estimated using a maximum likelihood framework to account for unphased mutations, whereas we use a normalization approach. The close similarity between our results, however, suggests that the effects of this difference are minor, and we therefore continued to use the Gao et al. parameters in further comparisons.

We tested whether our inferred sex specific baboon DNM counts were consistent with a male-to-female mutation ratio, *α*, that is 2.2-fold lower than that reported in humans. For the purposes of this test, we defined *α* as the ratio of paternal to maternal mutations at typical ages of reproduction (see section below for discussion on reproductive ages). Using the Gao et al. three-generation age effect estimates yielded a human *α* value of 4.106 (27). We thus formally performed a likelihood ratio test on the null hypothesis that in baboons *α* = 4.106/2.2 = 1.87. We used the mle2 function from the bbmle R package to compute maximum likelihood estimates (MLEs) using the BFGS optimization method and tested a range of initial values for *α* (under the free *α* model).

We tested whether the age effect measured in baboon fathers was consistent with a model in which mutations track cell divisions. Under this model we would expect the paternal age effect in baboons, 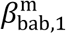, to be 16/11 greater than that measured in humans, reflecting the ratio of their cell division rates (see below). Using the estimates from Gao et al. yielded the null hypothesis 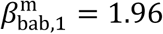. We similarly tested whether the baboon paternal age effect was equal to that of humans (null hypothesis 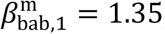). For both tests, we calculated the MLE under the fixed 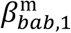 model using mle2 and a range of initial values for the intercept, 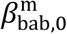. The MLE outputted by the glm R function in our initial age effect regression served as the likelihood estimate under the free 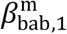 model.

### Testing an Exponential Maternal Age Effect Model

Gao et al. showed an exponential age effect model to be a slightly superior fit to the maternal deCODE data than a linear model, a result apparently driven by mothers above the age of 40 (27). Since mothers in our sample tend to be older than typical, we tested an exponential model as well. To this end, we calculated the Akaike information criterion for our human maternal data under both the linear model of Gao et al. as well as their exponential model (using parameter estimates obtained from three-generation families), applying the same genome scaling as before to both models.

### Estimating the Male-to-Female Mutation Ratio

In order to reliably compare the male-to-female mutation ratios, *α*, between our species, we considered only those DNMs that were transmitted (i.e., observed at least once in an F_2_), reasoning that this criterion should eliminate most false positive DNMs, which would otherwise bias our estimate downwards. We estimated *α* as the ratio of paternally-to maternally-phased DNMs using the phasing results described above (see Fig. 3A). For each species, we quantified the uncertainty in our estimate by segmenting the genome into blocks of a chosen size, bootstrap resampling these blocks 1000 times, re-calculating the male-to-female DNM ratio for each replicate, and then taking the 2.5–97.5% quantile range across the replicates as our 95% confidence interval. We used blocks of 50 cM width in humans, based on the HapMap Phase II b37 genetic map (73,74), and 50 Mb blocks in baboons. We performed a chi-squared statistical test on the observed distribution of paternal and maternal DNM counts between the two species using the chisq.test function in R.

### Comparing Mammalian Estimates of the Male-to-Female Mutation Ratio

We collected estimates of the male-to-female mutation ratio, *α*, from published mammalian pedigree sequencing studies containing phased DNMs (Table 1, Fig. 3B). When possible, we used the unaltered *α* estimates reported by the authors. If a ratio was not reported, we calculated an *α* estimate using the ratio of the total number of DNMs phased to the male versus the female genome. For the Jónsson et al. (16) human estimate, we included only pedigree-phased DNMs from the 225 three-generation families. For each study, we also calculated the mean of the reported ages of the father at conception, excluding fathers if their F_1_ offspring did not contribute phased DNMs to the *α* estimate. Since parental ages were unavailable in the cattle study, we relied instead on typical ages from the literature (49,50). Note that since Besenbacher et al. reanalyzed the chimpanzee data generated by Venn et al., one of the earliest non-human pedigree sequencing studies, we chose to reference only the results of Besenbacher et al. (23,48). We performed a linear regression in R of the mammalian *α* estimates, including our own, against the mean paternal age of the study. To avoid duplicating species, estimates from Jónsson et al. (human) and Tatsumoto et al. (chimpanzee) were not included (16,47).

### Comparing Mutation Spectra

We examined the single base pair point mutation types of the transmitted DNMs, using the species’ reference genome to categorize C>T transitions into those that occurred within (CpG>TpG) and outside of a CpG context (CpH>TpH), for a total of seven mutation types. To obtain a baseline expectation for “mutation spectrum” (i.e., the proportion of each mutation type), we compared our data to human DNMs published by Jónsson et al. as well as low frequency polymorphisms in the sample of 89 unrelated baboons described above (16,53). We restricted our use of the Jónsson et al. dataset to the 14,629 autosomal mutations called from three-generation pedigrees, which is similar to our study design. For the unrelated baboons, we identified non-singleton, autosomal SNPs with minor allele frequencies less than 0.05 that were genotyped in all 89 individuals (11,818,769 SNPs total). These low frequency variants should reflect the broad patterns of the de novo mutation spectrum, as most will have arisen recently and been only minimally affected by selection and biased gene conversion (25,26,83). We categorized SNPs by type under the assumption that the major allele is ancestral. Uncertainty in the mutation type proportions was computed by bootstrap resampling either 50 cM blocks, in humans, or 50 Mb blocks, in baboons, 1000 times and re-computing the proportions for each replicate. We used the 2.5–97.5% quantile range across the replicates as our 95% confidence interval.

### Comparing Yearly Mutation Rates and Substitution rates

To calculate yearly mutation rates per base pair, *μ_y,x_*, we used the model of ref. (36), which accounts for sex differences in generation times by dividing the sex-averaged mutation rate by the sex-averaged generation times:

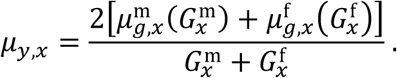

We note that here the generation times are the average over the lineage of species *x* and thus comparable (in principle) to yearly neutral substitution rates. For humans, we assumed average paternal and maternal generation times of 32.0 and 28.2 years, respectively; in baboons, we assumed an average generation time of 10.7 years in males and 10.2 years in females (see below for description of age choices).

The number of neutral substitutions, *K_x_*, arising in lineage *x* over the *T_x_* years since the mean time to the human-baboon common ancestor is *K_x_* = *μ_y,x_T_x_*. Focusing on neutral sites, we therefore expect that

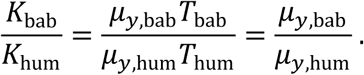

We used the substitution rates at putatively neutral sites in human and baboon lineages reported by ref. (21), for all mutation types. We used the randomly drawn regression coefficients described above to propagate uncertainty in the sex-specific mutation rate estimates on to the relative per-year mutation rate (but treating FDR and FNR as fixed at their estimated values). We plotted the human and baboon lineages with a marmoset (*Callithrix jacchus*) outgroup using ggtree in R and with branch lengths corresponding to the substitution rates reported by ref. (21) for all mutation types (84).

### Inferring Split Times of Humans and Baboons

We were interested in estimating the split time of humans (an ape) and baboons (an Old World Monkey). For humans and OWMs, the split time and the mean time to the common ancestor *T_x_* is relatively close, with an estimated difference of 2–5 My (42). The value *T_x_* can be estimated from substitution rates, given an estimate of the yearly mutation rate, but relies on sex-specific generation times along each lineage, which are unknown (36). We therefore explored the effects of life history trait values on inferred mean times to the common ancestor (*T_x =_ K_x_*/*μ_y,x_*) using a range of plausible values. To this end, we used the mean age of first reproduction of 14 New World monkey species (3.0 years) as the lower bound on paternal generation times for both species (45). For the upper bound on human paternal generation times, we used the modern generation times of agricultural humans (roughly 32.0 years); for baboons, we intentionally selected an upper bound of 15 years that is greater than the reported paternal generation times of modern baboon populations (10.7 years) in order to allow for the possibility of older than present-day generation times in the evolutionary history of the baboon lineage (see below for a discussion of generation times). We also varied the paternal-to-maternal generation time ratio from 0.8 to 1.2 (36). We estimated *T_x_* using the substitution rate estimates from ref. (21) as above along with the adjusted Gao et al. values for the intercept and age effect (described in “Testing Age Effects and Intercepts”).

## References

1. Feng C, Pettersson M, Lamichhaney S, Rubin CJ, Rafati N, Casini M, et al. Moderate nucleotide diversity in the Atlantic herring is associated with a low mutation rate. Elife. 2017;6.

2. Harland C, Charlier C, Karim L, Cambisano N, Deckers M, Mullaart E, et al. Frequency of mosaicism points towards mutation-prone early cleavage cell divisions. bioRxiv [Internet]. 2016;1–22.

3. Goldberg ME, Harris K. Great ape mutation spectra vary across the phylogeny and the genome due to distinct mutational processes that evolve at different rates. bioRxiv [Internet]. 2019 Jan 1;805598.

4. Lindsay SJ, Rahbari R, Kaplanis J, Keane T, Hurles ME. Similarities and differences in patterns of germline mutation between mice and humans. Nat Commun [Internet]. 2019 Dec 6;10(1):4053.

5. Harris K. Evidence for recent, population-specific evolution of the human mutation rate. Proc Natl Acad Sci U S A [Internet]. 2015;112(11):3439–44.

6. Harris K, Pritchard JK. Rapid evolution of the human mutation spectrum. Elife. 2017;6:1–17.

7. Mallick S, Li H, Lipson M, Mathieson I, Gymrek M, Racimo F, et al. The Simons Genome Diversity Project: 300 genomes from 142 diverse populations. Nature [Internet]. 2016 Oct 21;538(7624):201–6.

8. Narasimhan VM, Rahbari R, Scally A, Wuster A, Mason D, Xue Y, et al. Estimating the human mutation rate from autozygous segments reveals population differences in human mutational processes. Nat Commun [Internet]. 2017;8(1):303.

9. Ruan Y, Chen B, Chen Q, Wen H, Wu C-I. Mutations beget more mutations – The baseline mutation rate and runaway accumulation. bioRxiv [Internet]. 2019 Jan 1;690099.

10. Sung W, Ackerman MS, Miller SF, Doak TG, Lynch M. Drift-barrier hypothesis and mutation-rate evolution. Proc Natl Acad Sci [Internet]. 2012;109(45):18488–92.

11. Goodman M. Rates of molecular evolution: The hominoid slowdown. BioEssays [Internet]. 1985 Jul;3(1):9–14.

12. Wu CI, Li WH. Evidence for higher rates of nucleotide substitution in rodents than in man. Proc Natl Acad Sci [Internet]. 1985 Mar 1;82(6):1741–5.

13. Wilson MA, Makova KD. Evolution and Survival on Eutherian Sex Chromosomes. Gojobori T, editor. PLoS Genet [Internet]. 2009 Jul 17;5(7):e1000568.

14. Ségurel L, Wyman MJ, Przeworski M, Laure S, Wyman MJ, Przeworski M. Determinants of Mutation Rate Variation in the Human Germline. Annu Rev Genomics Hum Genet [Internet]. 2014;(May):1–24.

15. Gao Z, Wyman MJ, Sella G, Przeworski M. Interpreting the Dependence of Mutation Rates on Age and Time. PLoS Biol. 2016;14(1):1–16.

16. Jónsson H, Sulem P, Kehr B, Kristmundsdottir S, Zink F, Hjartarson E, et al. Parental influence on human germline de novo mutations in 1,548 trios from Iceland. Nature [Internet]. 2017 Sep;549(7673):519.

17. Wong WSW, Solomon BD, Bodian DL, Kothiyal P, Eley G, Huddleston KC, et al. New observations on maternal age effect on germline de novo mutations. Nat Commun [Internet]. 2016;7(10486):1–10.

18. Goldmann JM, Wong WSW, Pinelli M, Farrah T, Bodian D, Stittrich AB, et al. Parent-of-origin-specific signatures of de novo mutations. Nat Genet [Internet]. 2016;48(8):935–9.

19. Koch EM, Schweizer RM, Schweizer TM, Stahler DR, Smith DW, Wayne RK, et al. De novo mutation rate estimation in wolves of known pedigree. Mol Biol Evol. 2019 Jul 12;36(11):2536–47.

20. Yi S, Ellsworth DL, Li W-H. Slow Molecular Clocks in Old World Monkeys, Apes, and Humans. Mol Biol Evol [Internet]. 2002 Dec 1;19(12):2191–8.

21. Moorjani P, Amorim CEG, Arndt PF, Przeworski M. Variation in the molecular clock of primates. Proc Natl Acad Sci. 2016;(15):1–39.

22. Thomas GWC, Wang RJ, Puri A, Harris RA, Raveendran M, Hughes DST, et al. Reproductive longevity predicts mutation rates in primates. Curr Biol. 2018;28(19):3193–7.

23. Besenbacher S, Hvilsom C, Marques-Bonet T, Mailund T, Schierup MH. Direct estimation of mutations in great apes reconciles phylogenetic dating. Nat Ecol Evol [Internet]. 2019 Feb 21;3(2):286–92.

24. Wang RJ, Thomas GWC, Raveendran M, Harris RA, Doddapaneni H, Muzny DM, et al. Paternal age in rhesus macaques is positively associated with germline mutation accumulation but not with measures of offspring sociability. bioRxiv [Internet]. 2019 Jan 1:706705.

25. Rahbari R, Wuster A, Lindsay SJ, Hardwick RJ, Alexandrov LB, Al Turki S, et al. Timing, rates and spectra of human germline mutation. Nat Genet [Internet]. 2015;48(December):1–11.

26. Agarwal I, Przeworski M. Signatures of replication timing, recombination, and sex in the spectrum of rare variants on the human X chromosome and autosomes. Proc Natl Acad Sci [Internet]. 2019 Sep 3;116(36):17916–24.

27. Gao Z, Moorjani P, Sasani TA, Pedersen BS, Quinlan AR, Jorde LB, et al. Overlooked roles of DNA damage and maternal age in generating human germline mutations. Proc Natl Acad Sci. 2019;201901259.

28. Onyango PO, Gesquiere LR, Altmann J, Alberts SC. Puberty and dispersal in a wild primate population. Horm Behav. 2013;64(2):240–9.

29. Chowdhury AK, Steinberger E. A study of germ cell morphology and duration of spermatogenic cycle in the baboon, Papio anubis. Anat Rec [Internet]. 1976 Jun;185(2):155–69.

30. Nielsen CT, Skakkebæk NE, Richardson DW, Darling JAB, Hunter WM, Jørgensen M, et al. Onset of the Release of Spermatozoa (Spermarche) in Boys in Relation to Age, Testicular Growth, Pubic Hair, and Height*. J Clin Endocrinol Metab [Internet]. 1986;62(3):532–5.

31. Helgason A, Hrafnkelsson B, Gulcher JR, Ward R, Stefánsson K. A Populationwide Coalescent Analysis of Icelandic Matrilineal and Patrilineal Genealogies: Evidence for a Faster Evolutionary Rate of mtDNA Lineages than Y Chromosomes. Am J Hum Genet [Internet]. 2003 Jun;72(6):1370–88.

32. Heller CG, Clermont Y. Spermatogenesis in man: an estimate of its duration. Science (80-). 1963;140(3563):184–6.

33. Drost JB, Lee WR. Biological basis of germline mutation: comparisons of spontaneous germline mutation rates among drosophila, mouse, and human. Environ Mol Mutagen. 1995;25(S2):48–64.

34. Moorjani P, Gao Z, Przeworski M. Human germline mutation and the erratic evolutionary clock. PLoS Biol [Internet]. 2016;14(10).

35. Van der Auwera GA, Carneiro MO, Hartl C, Poplin R, del Angel G, Levy-Moonshine A, et al. From FastQ Data to High-Confidence Variant Calls: The Genome Analysis Toolkit Best Practices Pipeline. In: Current Protocols in Bioinformatics [Internet]. Hoboken, NJ, USA: John Wiley & Sons, Inc.; 2013. p. 11.10.1–11.10.33.

36. Amster G, Sella G. Life history effects on the molecular clock of autosomes and sex chromosomes. Proc Natl Acad Sci U S A [Internet]. 2016;113(6):1588–93.

37. Kong A, Frigge ML, Masson G, Besenbacher S, Sulem P, Magnusson G, et al. Rate of de novo mutations and the importance of father’s age to disease risk. Nature [Internet]. 2012;488(7412):471–5.

38. Pfeifer SP. Direct estimate of the spontaneous germ line mutation rate in African green monkeys. Evolution (N Y) [Internet]. 2017 Dec;71(12):2858–70.

39. Goldmann JM, Veltman JA, Gilissen C. De Novo Mutations Reflect Development and Aging of the Human Germline. Trends Genet [Internet]. 2019 Oct 11.

40. Scally A. Mutation rates and the evolution of germline structure. Philos Trans R Soc B Biol Sci [Internet]. 2016 Jul 19;371(1699):20150137.

41. Martin AP, Palumbi SR. Body size, metabolic rate, generation time, and the molecular clock. Proc Natl Acad Sci [Internet]. 1993;90(9):4087–91.

42. Steiper ME, Young NM, Sukarna TY. Genomic data support the hominoid slowdown and an Early Oligocene estimate for the hominoid-cercopithecoid divergence. Proc Natl Acad Sci U S A [Internet]. 2004;101(49):17021–6.

43. Seiffert ER, Simons EL, Fleagle JG, Godinot M. Paleogene Anthropoids. In: Cenozoic Mammals of Africa. Univ of California Press; 2012. p. 369–92.

44. Harrison T. Catarrhine origins. A companion to paleoanthropology. 2013;376–96.

45. Wootton JT. The effects of body mass, phylogeny, habitat, and trophic level on mammalian age at first reproduction. Evolution (N Y). 1987;41(4):732–49.

46. Scally A, Durbin R. Revising the human mutation rate: implications for understanding human evolution. Nat Rev Genet [Internet]. 2012;13(10):745–53.

47. Tatsumoto S, Go Y, Fukuta K, Noguchi H, Hayakawa T, Tomonaga M, et al. Direct estimation of de novo mutation rates in a chimpanzee parent-offspring trio by ultra-deep whole genome sequencing. Sci Rep [Internet]. 2017;7(1):13561.

48. Venn O, Turner I, Mathieson I, Groot N De, Bontrop R, Mcvean G. Strong male bias drives germline mutation in chimpanzees. Science (80-). 2014;344(6189):1272–5.

49. Sørensen AC, Sørensen MK, Berg P. Inbreeding in Danish Dairy Cattle Breeds. J Dairy Sci [Internet]. 2005 May;88(5):1865–72.

50. Murray C, Huerta-Sanchez E, Casey F, Bradley DG. Cattle demographic history modelled from autosomal sequence variation. Philos Trans R Soc B Biol Sci [Internet]. 2010 Aug 27;365(1552):2531–9.

51. Hostetler JA, Opitz JM, Reynolds JF. History and relevance of the Hutterite population for genetic studies. Am J Med Genet [Internet]. 1985 Nov;22(3):453–62.

52. Kosova G, Scott NM, Niederberger C, Prins GS, Ober C. Genome-wide association study identifies candidate genes for male fertility traits in humans. Am J Hum Genet [Internet]. 2012 Jun 8;90(6):950–61.

53. Robinson JA, Belsare S, Birnbaum S, Newman DE, Chan J, Glenn JP, et al. Analysis of 100 high-coverage genomes from a pedigreed captive baboon colony. Genome Res. 2019;29(5):848–56.

54. Li H. seqtk [Internet]. Available from: https://github.com/lh3/seqtk

55. Li H. trimadap [Internet]. Available from: https://github.com/lh3/trimadap

56. Li H. Aligning sequence reads, clone sequences and assembly contigs with BWA-MEM. arXiv Prepr arXiv13033997. 2013;

57. Faust GG, Hall IM. SAMBLASTER: fast duplicate marking and structural variant read extraction. Bioinformatics. 2014;30(17):2503–5.

58. Li H, Handsaker B, Wysoker A, Fennell T, Ruan J, Homer N, et al. The sequence alignment/map format and SAMtools. Bioinformatics. 2009;25(16):2078–9.

59. McKenna A, Hanna M, Banks E, Sivachenko A, Cibulskis K, Kernytsky A, et al. The Genome Analysis Toolkit: a MapReduce framework for analyzing next-generation DNA sequencing data. Genome Res. 2010;20(9):1297–303.

60. Li H. A statistical framework for SNP calling, mutation discovery, association mapping and population genetical parameter estimation from sequencing data. Bioinformatics. 2011;27(21):2987–93.

61. DePristo MA, Banks E, Poplin R, Garimella KV, Maguire JR, Hartl C, et al. A framework for variation discovery and genotyping using next-generation DNA sequencing data. Nat Genet. 2011 May;43(5):491–8.

62. Quinlan AR, Hall IM. BEDTools: a flexible suite of utilities for comparing genomic features. Bioinformatics. 2010;26(6):841–2.

63. Livne OE, Han L, Alkorta-Aranburu G, Wentworth-Sheilds W, Abney M, Ober C, et al. PRIMAL: Fast and Accurate Pedigree-based Imputation from Sequence Data in a Founder Population. McHardy AC, editor. PLOS Comput Biol [Internet]. 2015 Mar 3;11(3):e1004139.

64. Zerbino DR, Achuthan P, Akanni W, Amode MR, Barrell D, Bhai J, et al. Ensembl 2018. Nucleic Acids Res [Internet]. 2018 Jan 4;46(D1):D754–61.

65. Paten B, Herrero J, Beal K, Fitzgerald S, Birney E. Enredo and Pecan: genome-wide mammalian consistency-based multiple alignment with paralogs. Genome Res. 2008;18(11):1814–28.

66. Earl D, Paten B, Diekhans M. mafTools [Internet]. Available from: https://github.com/dentearl/mafTools

67. Haeussler M, Zweig AS, Tyner C, Speir ML, Rosenbloom KR, Raney BJ, et al. The UCSC Genome Browser database: 2019 update. Nucleic Acids Res. 2018;47(D1):D853–D858.

68. Wall JD, Schlebusch SA, Alberts SC, Cox LA, Snyder-Mackler N, Nevonen KA, et al. Genomewide ancestry and divergence patterns from low-coverage sequencing data reveal a complex history of admixture in wild baboons. Mol Ecol [Internet]. 2016 Jul;25(14):3469–83.

69. Delaneau O, Marchini J, Zagury J-F. A linear complexity phasing method for thousands of genomes. Nat Methods [Internet]. 2011;9(2):179–81.

70. Delaneau O, Zagury J-F, Marchini J. Improved whole-chromosome phasing for disease and population genetic studies. Nat Methods [Internet]. 2012;10(1):5–6.

71. Purcell S, Chang C. PLINK 1.9.

72. Chang CC, Chow CC, Tellier LCAM, Vattikuti S, Purcell SM, Lee JJ. Second-generation PLINK: rising to the challenge of larger and richer datasets. Gigascience. 2015;4(1):7.

73. The International HapMap Consortium. A haplotype map of the human genome. Nature [Internet]. 2005 Oct;437(7063):1299–320.

74. The International HapMap Consortium. The International HapMap Project. Nature [Internet]. 2003 Dec;426(6968):789–96.

75. Cox LA, Mahaney MC, VandeBerg JL, Rogers J. A second-generation genetic linkage map of the baboon (Papio hamadryas) genome. Genomics. 2006;88(3):274–81.

76. Dumont BL, Payseur BA. Evolution of the genomic rate of recombination in mammals. Evolution (N Y). 2008 Feb;62(2):276–94.

77. O’Connell J, Gurdasani D, Delaneau O, Pirastu N, Ulivi S, Cocca M, et al. A General Approach for Haplotype Phasing across the Full Spectrum of Relatedness. PLoS Genet [Internet]. 2014;10(4):1–21.

78. Untergasser A, Cutcutache I, Koressaar T, Ye J, Faircloth BC, Remm M, et al. PRIMER3 [Internet]. Available from: http://sourceforge.net/projects/primer3/

79. Untergasser A, Cutcutache I, Koressaar T, Ye J, Faircloth BC, Remm M, et al. Primer3--new capabilities and interfaces. Nucleic Acids Res [Internet]. 2012/06/21. 2012 Aug;40(15):e115–e115.

80. Ye J, Coulouris G, Zaretskaya I, Cutcutache I, Rozen S, Madden TL. Primer-BLAST: a tool to design target-specific primers for polymerase chain reaction. BMC Bioinformatics [Internet]. 2012 Jun 18;13:134.

81. GSL Biotech. SnapGene.

82. Tacutu R, Craig T, Budovsky A, Wuttke D, Lehmann G, Taranukha D, et al. Human Ageing Genomic Resources: integrated databases and tools for the biology and genetics of ageing. Nucleic Acids Res. 2012;41(D1):D1027–D1033.

83. Carlson J, Locke AE, Flickinger M, Zawistowski M, Levy S, Myers RM, et al. Extremely rare variants reveal patterns of germline mutation rate heterogeneity in humans. Nat Commun. 2018;9(1):3753.

84. Yu G, Smith DK, Zhu H, Guan Y, Lam TT-Y. ggtree: an R package for visualization and annotation of phylogenetic trees with their covariates and other associated data. Methods Ecol Evol. 2017;8(1):28–36.

