## Supplementary Information for "A comparison of humans and baboons suggests germline mutation rates do not track cell divisions"

**Supplementary Figures**

Figure S1. Size distributions and transmission status of clusters of DNM calls.
Figure S2. Estimates of false negative and false discovery rates of DNM calls.
Figure S3. Pedigrees with sample identifiers.
Figure S4. Sanger sequencing validation results and chromatogram examples.
Figure S5. Consistency of mutation rates within and outside of the orthologous regions.
Figure S6. Human and baboon mutation and substitution rates per year, distinguishing types subject and not subject to GC-biased gene conversion.
Figure S7. Estimated false discovery rates against F1 depth of coverage, across baboon trios.

**Supplementary Tables**

Table S1. DNA primers and results of Sanger sequencing in H14.
Table S2. Age, sex, and pedigree relationships of sequenced humans.
Table S3. Age, sex, and pedigree relationships of sequenced baboons.
Table S4. List of de novo mutation filters.
Table S5. Life history traits of humans and baboons.
Supplementary Data. Putative de novo mutations in humans and baboons. Coverage, false negative rates, false discovery rates, and callable genome sizes of human and baboon trios.

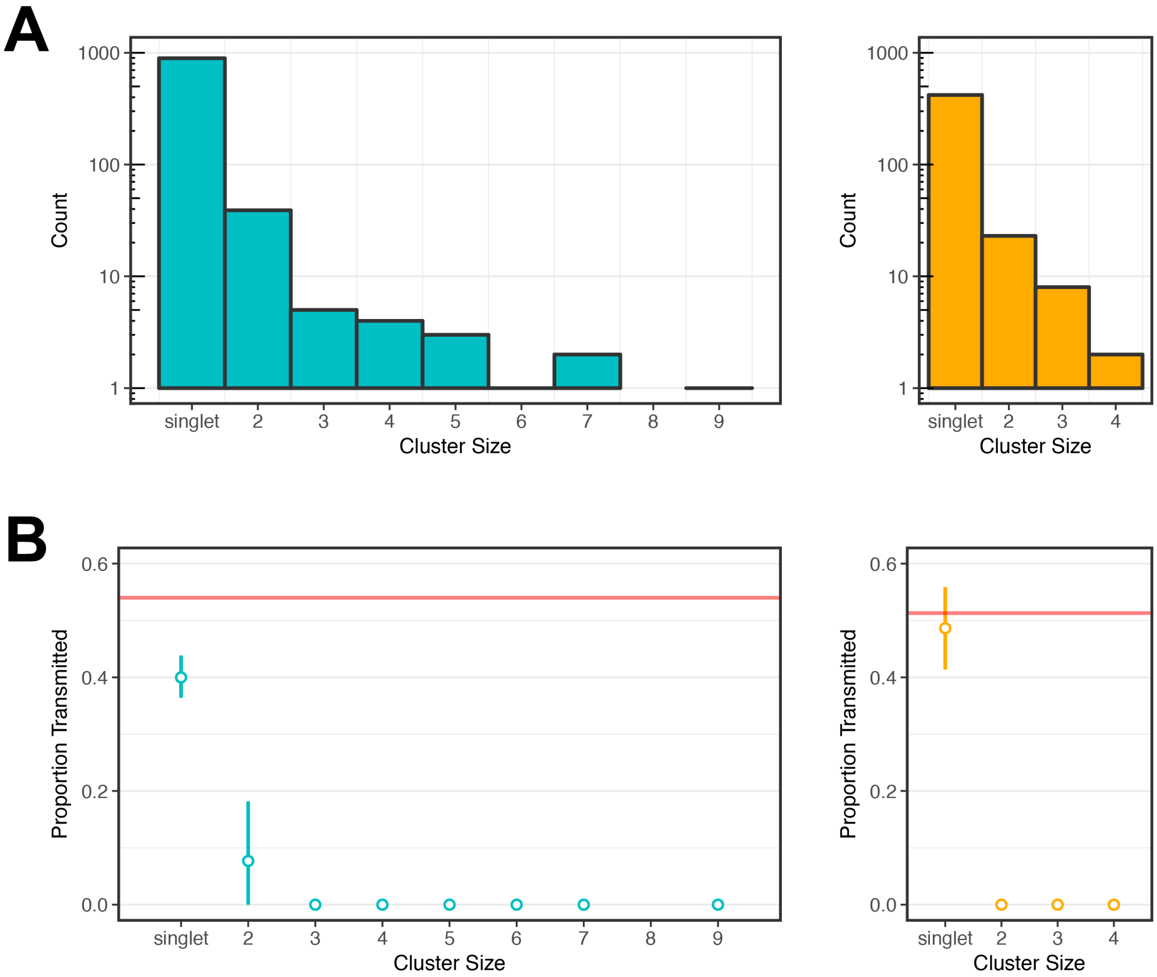


**Figure S1. Size distributions and transmission status of clusters of DNM calls.** (A) Distribution of DNM cluster sizes (defined as the number of DNM calls within 100 bp of each other) in humans (left, teal) and baboons (right, orange). The height of each bar denotes the number of DNM clusters of the given size, as labeled on the x-axis. (B) Proportion of DNM clusters observed transmitted to the $F_2$ generation, by size, in humans (left, teal) and baboons (right, orange). For each species, the horizontal red line denotes the mean proportion of DNMs expected to be observed in the $F_2$ generation if called in the F1, as determined from simulations (see Methods). Vertical lines denote the 95% confidence interval, as determined by bootstrap resampling clusters by 50 cM blocks in humans and 50 Mb blocks in baboons.

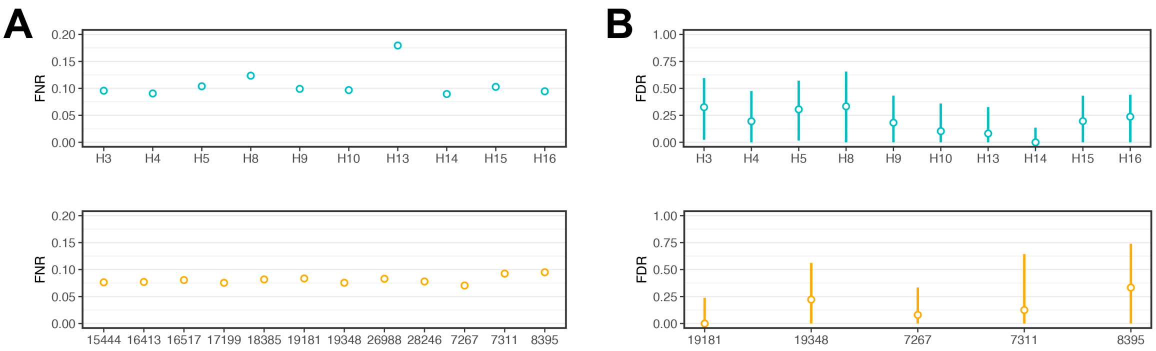


**Figure S2. Estimates of false negative and false discovery rates of DNM calls.** False negative rates (in A) and false discovery rates (in B) of the DNM-calling pipeline estimated for each trio across humans (top, teal) and baboons (bottom, orange). In all plots, sample identifiers of the focal $F_1$ are provided as labels on the x-axis. False negative rates were estimated by applying the filtering pipeline to a set of simulated DNMs (20,000 per trio). For false negative rates, 95% confidence intervals from bootstrap resampling of mutations are narrow and hidden by the points. False discovery rates were inferred using a transmission-based approach in which the proportion of DNMs called in the $F_1$ that were also observed in the $F_2$ was compared against an expected proportion determined from simulations. Only the five baboon $F_1$ individuals with at least one sequenced $F_2$ offspring are depicted. Vertical lines denote 95% confidence intervals from bootstrap resampling of 50 cM blocks in humans (1,2) and 50 Mb blocks in baboons. See Methods for details on error rate estimation.

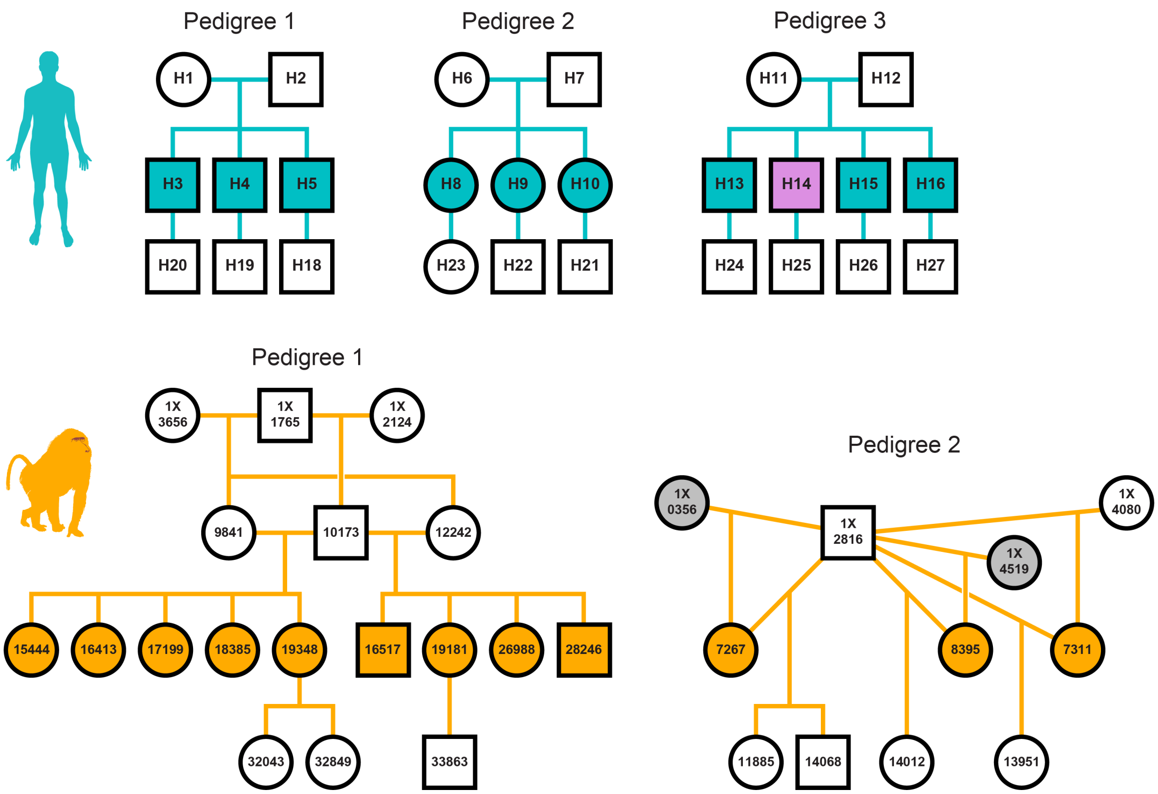


**Figure S3. Pedigrees with sample identifiers.** Reproduction of Figure 1 with unique sample identifiers for each individual in the human (top row, teal) and baboon (bottom row, orange) pedigrees. Sanger sequencing validation experiments focused on putative DNMs called in $F_1$ individual H14, highlighted in lavender. Baboon mothers 1X0356 and 1X4519, highlighted in grey, lacked birth dates and were thus excluded from age effect analyses.

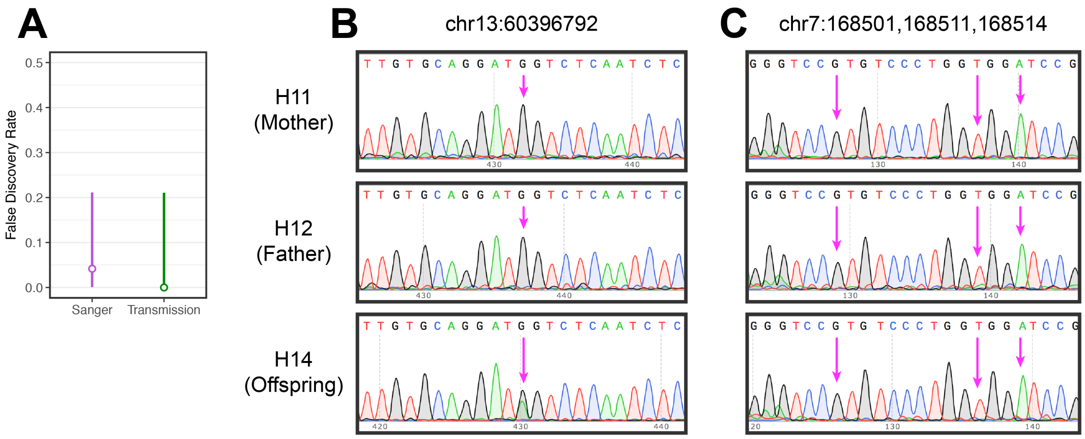


**Figure S4. Sanger sequencing validation results and chromatogram examples.** (A) Comparison of two methods for inferring the false discovery rate of DNMs in human $F_1$ individual H14. In one, the false discovery rate was inferred by Sanger resequencing of putative DNMs (purple) and in the second, by a transmission-based approach (green) in which the number of DNMs observed in the $F_2$ offspring of H14 was compared to the expectation determined by simulations (see main text). Vertical lines indicate the 95% confidence interval of the point estimates. For the Sanger data, the binomial confidence interval is indicated; for the transmission-based method, bootstrap resampling of 50 cM blocks (1,2) was used to obtain the interval. Examples of four-color chromatograms from Sanger resequencing of a genuine G/C>A/T singlet DNM (in B) and a cluster of three spurious DNMs (in C). Chromatograms for the mother (top row) and father (middle row) of H14 (bottom row) are provided. Magenta arrows indicate the positions of putative DNMs originally called from Illumina sequence alignments. Absolute positions of the DNMs are also given at the top of each column using hs37d5 reference genome coordinates. Note the overlapping G (black) and A (green) peaks at the genuine DNM site in H14, indicating a heterozygous AG genotype, and the absence of a similar signal at the clustered calls.

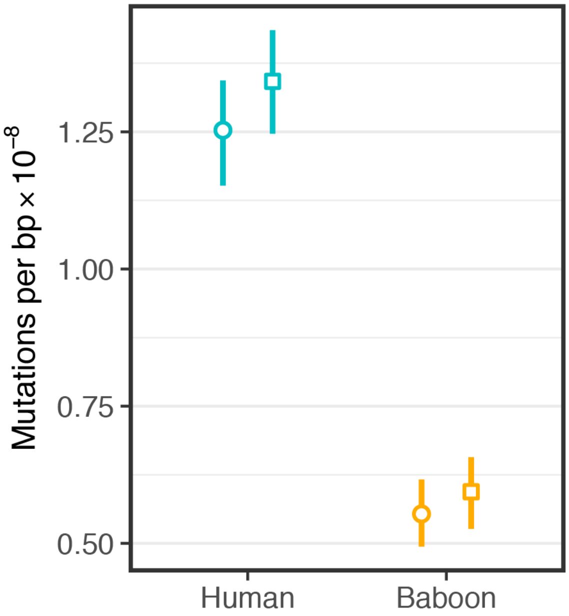


**Figure S5. Consistency of mutation rates within and outside of the orthologous regions.**

Mutation rates per generation were calculated for humans (teal) and baboons (orange). Circles denote rates calculated using mutations identified within regions of the genome identified as orthologous between the two species whereas squares represent estimates calculated using all callable regions of the genome. Considering all callable regions resulted in a slightly higher mutation rate in both species. This slight difference is expected since the orthologous regions are likely to be somewhat more conserved (due to natural selection).

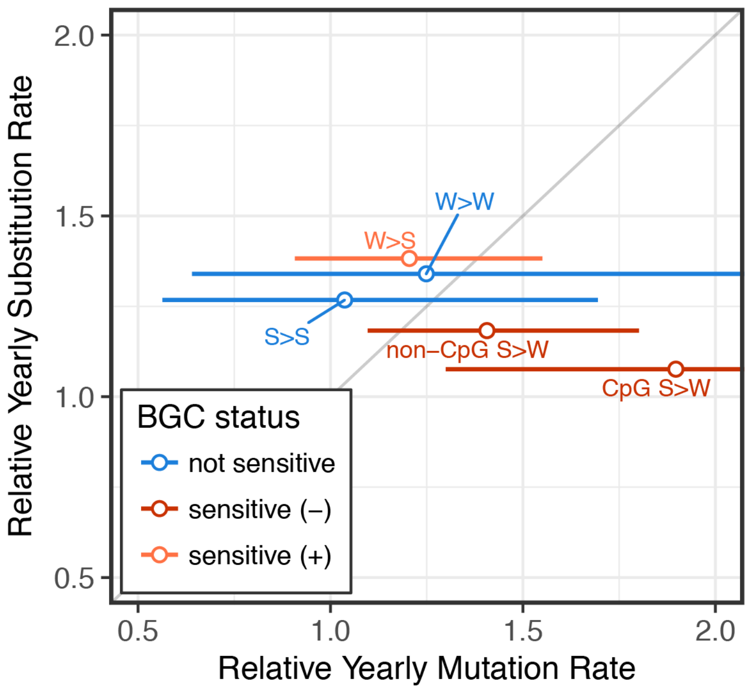


**Figure S6. Human and baboon mutation and substitution rates per year, distinguishing types subject and not subject to GC-biased gene conversion.** The ratio of yearly mutation and substitution rates in baboon relative to human, as estimated for the different possible types involving combinations of strong (S: G/C) and weak (W: A/T) base pairs. Each point denotes a different type, and strong-to-weak (S>W) types were separated into those that occurred at a CpG or a non-CpG site. Points are colored according to whether GC-biased gene conversion is expected to favor (light red), disfavor (dark red), or have no effect (blue) on the mutation type. The mutation rates used for these ratios (x-axis) were calculated in each species by aggregating information across all trios: for each type, we estimated the yearly mutation rate by collating the mutations and mutational opportunities of a given mutation type across $F_1$s, weighted by the error rates estimated for each $F_1$, and given the mean ages of reproduction of mothers and fathers. Baboon $F_1$ individuals 7267 and 8395 were excluded due to incomplete $P_0$ age information. Horizontal lines denote 95% confidence intervals on the mutation rate ratio computed by resampling within each individual using 50 cM blocks in humans and 50 Mb blocks in baboons. Upper confidence intervals for CpG S>W and W>W types extend out of frame to 2.8 and 2.1, respectively. Point estimates for the substitution rates in baboons and humans were taken from ref. (3). The identity line is drawn in grey for reference.

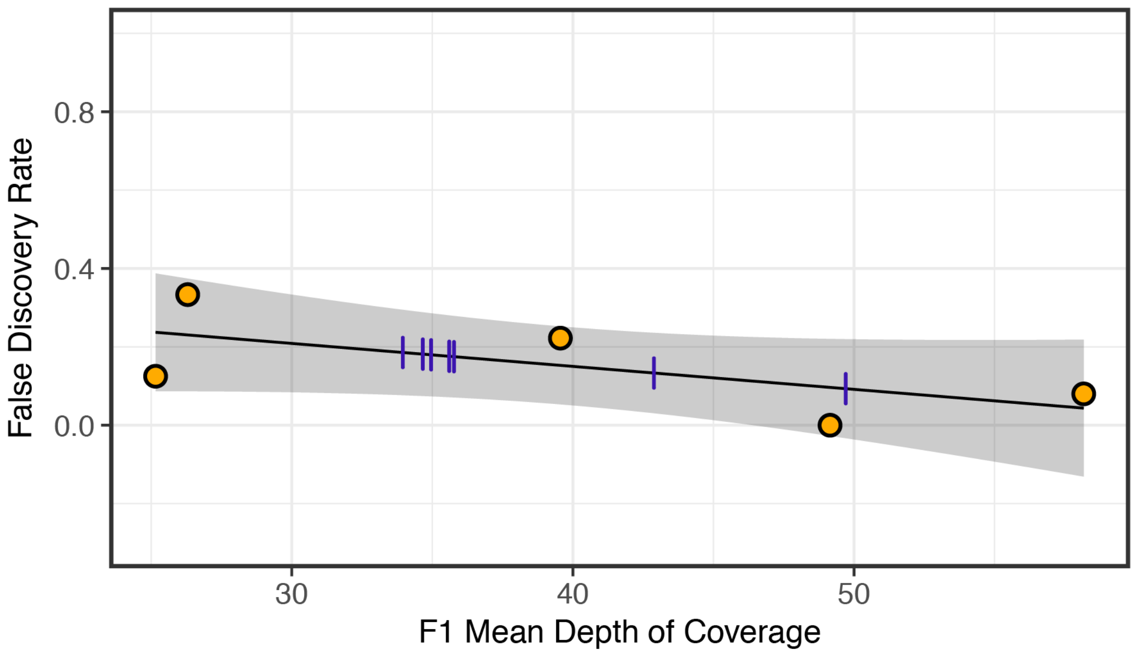


**Figure S7. Estimated false discovery rates against F1 depth of coverage, across baboon trios.** The false discovery rate (FDR) of DNM calls was inferred using a transmission-based approach in the five baboon $F_1$ individuals (orange points) for which an $F_2$ offspring was available. The black line indicates the best fit of a linear regression of these estimated FDRs against the mean depth of sequencing coverage in the same individuals. The grey shaded region denotes the 95% confidence intervals of the intercept and slope of the regression. The slope of the relationship is slightly negative (-0.0059), but not significantly so (p-value = 0.24). The regression fit was nonetheless used to predict FDRs in the seven baboon $F_1$s that lacked an $F_2$ generation from their mean depth of coverage. Purple hash lines on the fitted line mark the predicted FDRs of these seven baboon $F_1$ individuals. As expected from the insignificant slope, using the mean FDR across all individuals instead does not change the qualitative conclusions and yields a very similar estimate of the mutation rate in baboons: $5.55\times10^{-9}$ instead of $5.54\times10^{-9}$ per generation.

**Table S2. Age, sex, and pedigree relationships of sequenced humans.**

| ID | Birth Year | Sex | Pedigree | Father | Mother | Father Age at Birth (in days) | Mother Age at Birth (in days) | Mean Depth of Coverage |
|---|---|---|---|---|---|---|---|---|
| H1 | 1923 | F | 1 | . | . | . | . | 28.7 |
| H2 | 1923 | M | 1 | . | . | . | . | 26.8 |
| H3 | 1947 | M | 1 | H2 | H1 | 9063 | 9039 | 22.9 |
| H4 | 1954 | M | 1 | H2 | H1 | 11353 | 11329 | 24.6 |
| H5 | 1968 | M | 1 | H2 | H1 | 16717 | 16693 | 21.2 |
| H6 | 1919 | F | 2 | . | . | . | . | 31.1 |
| H7 | 1920 | M | 2 | . | . | . | . | 30.6 |
| H8 | 1940 | F | 2 | H7 | H6 | 7639 | 7767 | 20.1 |
| H9 | 1952 | F | 2 | H7 | H6 | 11876 | 12004 | 21.5 |
| H10 | 1962 | F | 2 | H7 | H6 | 15532 | 15660 | 20.1 |
| H11 | 1931 | F | 3 | . | . | . | . | 31.7 |
| H12 | 1923 | M | 3 | . | . | . | . | 30.7 |
| H13 | 1955 | M | 3 | H12 | H11 | 11640 | 8794 | 18.6 |
| H14 | 1959 | M | 3 | H12 | H11 | 13005 | 10159 | 22.8 |
| H15 | 1961 | M | 3 | H12 | H11 | 13740 | 10894 | 20.9 |
| H16 | 1971 | M | 3 | H12 | H11 | 17403 | 14557 | 22.0 |
| H18 | 2001 | M | 1 | H5 | . | 11780 | . | 24.3 |
| H19 | 1976 | M | 1 | H4 | . | 7975 | . | 23.9 |
| H20 | 1970 | M | 1 | H3 | . | 8158 | . | 22.6 |
| H21 | 1992 | F | 2 | . | H10 | . | 10934 | 25.3 |
| H22 | 1982 | M | 2 | . | H9 | . | 10851 | 24.2 |
| H23 | 1968 | M | 2 | . | H8 | . | 10106 | 24.5 |
| H24 | 1984 | M | 3 | H16 | . | 4635 | . | 23.1 |
| H25 | 1991 | M | 3 | H15 | . | 10952 | . | 23.7 |
| H26 | 1992 | M | 3 | H14 | . | 12284 | . | 25.0 |
| H27 | 1993 | M | 3 | H13 | . | 13979 | . | 22.9 |

**Table S3. Age, sex, and pedigree relationships of sequenced baboons.**

| ID | Birth Date (yyyy-mm-dd) | Sex | Pedigree | Father | Mother | Father Age at Birth (in days) | Mother Age at Birth (in days) | Mean Depth of Coverage |
|---|---|---|---|---|---|---|---|---|
| 10173 | 1991-02-23 | M | 1 | 1X1765 | 1X2124 | . | 4712 | 89.2 |
| 12242 | 1994-09-14 | F | 1 | 1X1765 | 1X3656 | . | 5260 | 34.6 |
| 9841 | 1990-08-17 | F | 1 | 1X1765 | 1X3656 | . | 3771 | 45.6 |
| 16517 | 2000-08-27 | M | 1 | 10173 | 12242 | 3473 | 2174 | 34.7 |
| 19181 | 2003-05-21 | F | 1 | 10173 | 12242 | 4470 | 3171 | 49.1 |
| 26988 | 2005-08-04 | F | 1 | 10173 | 12242 | 5276 | 3977 | 42.9 |
| 28246 | 2006-08-03 | M | 1 | 10173 | 12242 | 5640 | 4341 | 35.6 |
| 15444 | 1999-05-03 | F | 1 | 10173 | 9841 | 2991 | 3181 | 35.0 |
| 16413 | 2000-07-07 | F | 1 | 10173 | 9841 | 3422 | 3612 | 34.0 |
| 17199 | 2001-07-27 | F | 1 | 10173 | 9841 | 3807 | 3997 | 35.8 |
| 18385 | 2002-05-30 | F | 1 | 10173 | 9841 | 4114 | 4304 | 49.7 |
| 19348 | 2003-08-08 | F | 1 | 10173 | 9841 | 4549 | 4739 | 39.6 |
| 32043 | 2012-09-08 | F | 1 | . | 19348 | . | 3319 | 40.2 |
| 32849 | 2013-08-16 | F | 1 | . | 19348 | . | 3661 | 40.1 |
| 33863 | 2014-12-17 | M | 1 | . | 19181 | . | 4228 | 40.0 |
| 1X3656 | 1980-04-20 | F | 1 | . | . | . | . | 68.1 |
| 1X1765 | . | M | 1 | . | . | . | . | 62.7 |
| 1X2124 | 1978-03-31 | F | 1 | . | . | . | . | 69.3 |
| 1X2816 | 1979-02-13 | M | 2 | . | . | . | . | 78.7 |
| 1X0356 | . | F | 2 | . | . | . | . | 65.3 |
| 1X4519 | . | F | 2 | . | . | . | . | 26.2 |
| 1X4080 | 1981-10-14 | F | 2 | . | . | . | . | 26.7 |
| 7267 | 1987-01-18 | F | 2 | 1X2816 | 1X0356 | 2896 | . | 58.2 |
| 8395 | 1988-08-26 | F | 2 | 1X2816 | 1X4519 | 3482 | . | 26.3 |
| 7311 | 1987-02-10 | F | 2 | 1X2816 | 1X4080 | 2919 | 1945 | 25.2 |
| 11885 | 1994-04-11 | F | 2 | 1X2816 | 7267 | 5536 | 2640 | 39.6 |
| 14068 | 1997-09-23 | M | 2 | 1X2816 | 7267 | 6797 | 3901 | 69.6 |
| 14012 | 1997-08-23 | F | 2 | 1X2816 | 8395 | 6766 | 3284 | 27.1 |
| 13951 | 1997-07-03 | F | 2 | 1X2816 | 7311 | 6715 | 3796 | 22.7 |

**Table S4. List of de novo mutation filters.**

| Filter Name | Filter Description |
|---|---|
| Read depth | Only considered regions of the genome with depth of coverage passing a two-sided Poisson test with p-value > $2\times10^{-4}$ in all members of the trio, assuming that $\lambda$, the Poisson parameter, was equal to the mean read depth |
| Biallelic | Required exactly 2 alleles present in the sample cohort at the DNM site |
| Hard filters | QualByDepth (QD) > 2.0, FisherStrand (FS) < 60.0, RMSMappingQuality (MQ) > 40.0, MappingQualityRankSumTest (MQRankSum) > -12.5, ReadPosRankSumTest (ReadPosRankSum) > -8.0, StrandOddsRatio (SOR) < 3.0 |
| Variant quality | Required QUAL ≥ 100 at DNM site |
| Genotype quality | Required GQ > 40 in all three members of trio |
| Allelic balance in F1 | Performed a two-sided binomial test on allelic balance (proportion of reads supporting the DNM) under the null hypothesis that the true allelic balance in the F1 is 50%; removed DNMs with p-value ≤ 0.05 |
| Allelic depth in F1 | Required > 3 reads supporting DNM in child |
| Allelic depth in P0s | Required allelic depth (number of reads supporting the DNM) be 0 in at least one parent |
| Unrelated filter | Removed all putative DNMs that were previously observed in an unrelated individual (i.e., allele frequency > 0.05); this includes members of unrelated pedigrees. |

**Table S5. Life history traits of humans and baboons.**

| | Human | Baboon | References |
|---|---|---|---|
| Paternal age at reproduction (y) | 32.0 | 10.7 | (4; Jenny Tung, personal communication) |
| Maternal age at reproduction (y) | 28.2 | 10.2 | (4; Jenny Tung, personal communication) |
| Male age at puberty (y) | 13 | 5.41 | (5,6) |
| Sperm cycle length (d) | 16 | 11 | (7,8) |
| Gestation time (d) | 280 | 171 | (9) |

**References**

1. The International HapMap Consortium. A haplotype map of the human genome. Nature [Internet]. 2005 Oct;437(7063):1299–320. Available from: http://www.nature.com/articles/nature04226
2. The International HapMap Consortium. The International HapMap Project. Nature [Internet]. 2003 Dec;426(6968):789–96. Available from: https://doi.org/10.1038/nature02168
3. Moorjani P, Amorim CEG, Arndt PF, Przeworski M. Variation in the molecular clock of primates. Proc Natl Acad Sci. 2016;(15):1–39.
4. Jónsson H, Sulem P, Kehr B, Kristmundsdottir S, Zink F, Hjartarson E, et al. Parental influence on human germline de novo mutations in 1,548 trios from Iceland. Nature [Internet]. 2017; Available from: http://dx.doi.org/10.1038/nature24018
5. Nielsen CT, Skakkebæk NE, Richardson DW, Darling JAB, Hunter WM, Jørgensen M, et al. Onset of the Release of Spermatozoa (Spermarche) in Boys in Relation to Age, Testicular Growth, Pubic Hair, and Height*. J Clin Endocrinol Metab [Internet]. 1986;62(3):532–5. Available from: https://doi.org/10.1210/jcem-62-3-532
6. Onyango PO, Gesquiere LR, Altmann J, Alberts SC. Puberty and dispersal in a wild primate population. Horm Behav. 2013;64(2):240–9.
7. Heller CG, Clermont Y. Spermatogenesis in man: an estimate of its duration. Science (80- ). 1963;140(3563):184–6.
8. Chowdhury AK, Steinberger E. A study of germ cell morphology and duration of spermatogenic cycle in the baboon, Papio anubis. Anat Rec [Internet]. 1976 Jun;185(2):155–69. Available from: http://doi.wiley.com/10.1002/ar.1091850204
9. Tacutu R, Craig T, Budovsky A, Wuttke D, Lehmann G, Taranukha D, et al. Human Ageing Genomic Resources: integrated databases and tools for the biology and genetics of ageing. Nucleic Acids Res. 2012;41(D1):D1027--D1033.
